# Engineering a hybrid 3D construct for bone regeneration to promote simultaneous pre-vascularization and osteogenic differentiation in vitro

**DOI:** 10.64898/2026.05.06.723258

**Authors:** Sophia Dalfino, Sveva Fagiolino, Ivo Beeren, Marco Borrone, Francesco Alviano, Carlos Mota, Gianluca Tartaglia, Claudia Dolci, Lorenzo Moroni

## Abstract

Critical-sized bone defects represent a challenge in bone tissue engineering, due to insufficient vascularization that results in implant failure. Scaffold pre-vascularization is a promising strategy to create a functional microvascular network that integrates with host vasculature. In this study, we present a hybrid 3D construct comprising a hyaluronic acid-based hydrogel and a 3D printed polycaprolactone/β-tricalcium phosphate scaffold, to support vascular network formation and osteogenic differentiation. Peptide-functionalized (i.e. RGD, YIGSR, IKVAV, QK) hydrogels were obtained via thiol-ene chemistry, using two crosslinkers (PEG-diSH or MMP-diSH). Preliminary biological experiments assessed human mesenchymal stromal cells (hMSCs), endothelial cells (hUVECs), and their co-culture, on different gel formulations. All cell conditions displayed enhanced spreading and metabolic activity on gel formulations comprising RGD; thus these (i.e. RGD only and a combination of RGD/YIGSR) were selected for further studies. Cells were then mixed with the hydrogel precursor solutions, which were injected to embed the scaffolds and crosslinked using a UV lamp. After 7 days, tubule formation was observed only in co-culture conditions, highlighting the importance of cellular crosstalk for the formation of a vascular network. Significant differences were found across the tested formulations. In the RGD-PEG constructs, hUVECs formed tubule-like structures, surrounded by hMSCs, exhibiting pericyte-like behavior, supported by the upregulation of αSMA gene. Conversely, in the RGD/YIGSR-MMP conditions, hMSCs were mostly located on the scaffold fibers, and showed the highest expression of early osteogenic markers (RUNX2 and ALP). Overall, we demonstrated that the hybrid system with tailored hydrogel chemistry can support simultaneous microvascular organization and osteogenic commitment, offering a promising platform for bone tissue engineering applications. However, further studies involving longer culture periods will aim at clarifying the complex interplay between material composition, cell crosstalk and spatial organization and their influence on the maturation and stability of the vascular network.

## 1. Introduction

Mandibular bone defects arising from mandibular resections are nowadays repaired with fibula flap grafts, the clinical golden standard treatment [1]. However, many limitations are associated with this highly invasive intervention, such as limited availability, and donor site morbidity [2]. Bone tissue engineering (BTE) offers an alternative solution to repair such defects, based on synthetic grafts (i.e. scaffolds), which can be combined with cells or bioactive factors to guide tissue regeneration. Additive manufacturing (AM) techniques are commonly used to fabricate 3D scaffolds for mandibular applications, as they allow control over scaffold design and mechanical properties, and processability of broad range of materials [3]. Despite many advances have been made in enhancing bone formation by optimizing composite materials via the addition of inorganic components such as calcium phosphates [4,5], as well as the scaffold geometry [6,7], a major limitation in clinical translation remains lack of vascularization of the 3D constructs [8]. Blood vessel infiltration from the host tissue is slow (5-17 µm/h) and reaches only a depth of a few hundreds of micrometers [9,10], whereas tissue engineered constructs span typically from a few millimeters up to a centimeter in length and/or height. Without sufficient blood perfusion, from early stages after implantation onward, these 3D constructs cannot support bone formation. Therefore, their angiogenic properties need to be improved to promote vascularization throughout the 3D constructs.

Several approaches have been proposed to enhance angiogenic processes, such as transfecting host cells by inducing an overexpression of pro-angiogenic factors [11,12], or engineering materials that can release pro-angiogenic ions or growth factors over time [13]. However, these approaches usually rely on the body response in activating and recruiting cells, which typically results in slow vascularization [14]. Moreover, the angiogenic factors are usually released in a short time window, limiting their long-term effect, which may be associated to incomplete vascularization [15]. To overcome these issues, one interesting approach is *in vitro* pre-vascularization of 3D constructs, encapsulating co-cultures of human endothelial cells (hUVECs) and stromal cells, which allows the formation of a microvascular network prior implantation that can later inosculate with the host vasculature [16,17]. This approach could potentially offer a less invasive semi-autologous alternative to the fibula flap grafts for mandibular reconstruction. In addition, pre-vascularized engineered constructs may also serve as complementary solutions to autologous grafts in complex reconstructive cases, for instance by supporting vascular integration and tissue remodeling at the host-graft interfaces.

Hydrogel-based constructs are suitable candidates as 3D matrices that support such co-cultures, as they can mimic the natural extracellular matrix (ECM) environment in terms of mechanical and biological properties. Moreover, the hydrogel network can be modified with bioactive factors or peptides that can direct cell behavior [18]. Several naturally derived hydrogels such as alginate, gelatin, hyaluronic acid and collagen were reported in the context of vascularization for bone application [19,20]. Among them, hyaluronic acid (HA) represents a promising option since its natural abundance in human ECM. Importantly, HA is degradable by stromal cells, such as human mesenchymal stromal cells (hMSCs), which through hyaluronidase secretion can remodel the matrix and form cellular networks. Moreover, HA degradation products have been shown to possess pro-angiogenic and osteo-inductive properties [21–23]. HA can also interact with the CD44 surface receptor group of hMSCs promoting their proliferation and migration [24]. From a chemical perspective, the HA backbone can be easily modified with motives (e.g. RGD) for focal adhesion formation, enhancing its adhesive properties.

Hydrogels are typically too soft to be applied in a high loading environment such as bone [25]. Embedding them in 3D polymeric scaffolds reinforces their mechanical properties of multiple magnitudes, creating suitable hybrid constructs to promote both vascularization and osteogenic differentiation. For example, PLA scaffolds embedded in a GelMA-based hydrogel were shown to simultaneously mechanically reinforce the construct and to improve human adipose-derived stem cells osteogenic differentiation [26]. *Li et al.* presented a PCL-Maleimide electrospun scaffold covalently linked to a HA-SH hydrogel to co-culture ADC spheroids and endothelial cells to induce vascular-like network formation [27]. More recently, *Neves et al.* presented different seeding approaches of endothelial and stromal cells to induce vascular network formation in hybrid constructs made of a PEOT/PBT scaffold and an RGD-modified pectin hydrogel [28].

Most of the studies in the context of pre-vascularization primarily focus on hydrogel properties to support endothelial organization in the 3D constructs but often overlook the osteogenic potential of the synthetic support structure. In a previous study from our group, we designed composite scaffolds comprising of polycaprolactone (PCL)/β-tricalcium phosphate (TCP, 40% w/v), with a diamond shaped pore geometry, which successfully promoted osteogenesis in hMSCs [29]. In this study, we built upon this work by integrating a chemically modified HA-based hydrogel with the AM scaffold, to enable facile tunability of its biological properties. HA possesses both maleimide and norbornene units. This dual chemistry strategy allowed independent peptide functionalization and crosslinking, thus permitting controlled modulation of biochemical and mechanical properties. This modular approach introduced high degree of tunability, overcoming the limitations of traditional systems where both mechanisms occur on a single reactive group, reducing control over matrix design [30–32]. In our system, we selectively grafted bioactive peptides via thiol-Michael addition reactions and induced hydrogel formation using different crosslinkers (polyethylene glycol- and matrix metalloproteinase-based) via thiol-ene chemistry, respectively.

The scope of this study was to fundamentally investigate the role of different hydrogel chemistries, moving beyond the commonly used RGD only modifications [33–35], and their interplay with osteogenic cues from AM scaffolds on early endothelial organization and initial osteogenic commitment of human mesenchymal stromal cells (hMSCs). Accordingly, we performed short-term cell culture experiments to evaluate the early phases of cellular response. First, we assessed the effect of different hydrogel formulations on spreading and metabolic activity of hMSCs, hUVECs and their co-culture. We selected hMSCs and hUVECs as well-established cell models, combining the osteogenic and pericyte-like potential of hMSCs with the widely characterized hUVECs, which are considered a gold standard in tubule formation studies. The most promising cell-laden hydrogel formulations were then used in hybrid scaffolds to assess their pre-vascularization and osteogenic capacity through fluorescent imaging and gene expression. To the best of our knowledge no studies report the use of composite scaffolds with advanced geometries in combination with such a hydrogel.

## 2. Materials and methods

### 2.1 Materials

All materials were purchased from Merck, unless stated otherwise.

### 2.2 HA modification

HA-norbornene (Nb)-maleimide (Mal) was synthesized following a two-step reaction (**FigureA**), by adapting a previously establish procedure [36].

#### Hyaluronic acid-norbornene (HA-Nb)

In a typical reaction, sodium hyaluronate 200k (600 mg, 1.5 mmol, 1.0 equiv of carboxylic acid, Lifecore Biomedical, HA200K-5) was dissolved in 0.1 M 2-(N-morpholino) ethanesulfonic acid (MES) buffer (0.3M NaCl, pH 6.7), at 1% w/v. Then, 4-(4,6-Dimethoxytriazin-2-yl)-4-methylmorpholinium chloride (DMTMM, 830 mg, 3 mmol, 2 equiv) and 5-Methylamine-2-Norbornene (385 μL, 3 mmol, 2 equiv) were added to the solution. After stirring overnight, the product was precipitated in cold ethanol, washed with EtOH (3x), and washed with diethyl ether (3x). The material was re-dissolved in distilled H_2_O (*d*H_2_O), transferred to a 3kDa molecular weight cutoff (MWCO) Snakeskin dialysis membrane, and dialysis was performed against decreasing NaCl concentration in *d*H_2_O (from 50 mM to 0 mM, 3 water changes a day). The purified material was obtained as a white fluffy solid after freeze-drying and validated via ^1^H NMR. Final yield = 480 mg, 80%.

#### Hyaluronic acid – norbornene – maleimide (HA-Nb-Mal)

Subsequently, in a typical reaction, the lyophilized HA-Nb (440 mg, 1.1 mmol, 1.0 equiv of carboxylic acid groups) was dissolved in degassed MES buffer at 1% w/v. Then, DMTMM (304 mg, 1.1 mmol, 1 equiv) and 2-maleimidoethylamine trifluoroacetate salt (283 mg, 1.1 mmol, 1 equiv) were added. The solution was shortly purged with dry N_2_ gas and left stirring overnight. The product was purified via precipitation in cold ethanol, wash in cold EtOH (3x) and diethyl ether (3x). The product was re-dissolved in MilliQ water, transferred to a 3kDa molecular weight cutoff (MWCO) Snakeskin dialysis membrane and dialyzed against decreasing NaCl concentration in MQ (from 50 mM to 0 mM, 3 water changes a day). The purified material was obtained as a white fluffy solid after freeze-drying and validated via ^1^H NMR. Final yield = 313 mg, 72%.

### 2.3 Peptide functionalization

Different peptides were selected to be grafted on the polymer and their sequences are listed in **Table 1**. A schematic of the reaction is represented in **FigureB**. All peptides were obtained from Synpeptide Co., Ltd (Purity > 98%) and possessed a terminal cysteine terminal to enable coupling via thiol-Michael addition reaction to the polymer backbone. Before conjugation, the peptides were incubated in PBS containing a 10-fold molar excess of tris(2-carboxyethyl) phosphine (TCEP) for 1h at 37°C to remove the spontaneously formed disulfide bonds (5 mM peptide concentration) Then, 400 mL of the peptide solution(s), 1000 uL HA-Nb-Mal previously dissolved in PBS (4% w/v), and 600 mL of PBS was combined to obtain a 1:1 peptide – maleimide stoichiometry. The solution was left for 1h at RT. Excess of TECP was removed with ultrafiltration with a 3kDa MWCO (Amicon Ultra, Millipore) filters, according to manufacturer instructions.

**Table 1.** Peptides used to functionalize the Ha-Nb-Mal, via Thiol-Michael addition chemistry. 1.0 equivalent of peptides was added from a stock solution (5 mM), relative to Mal groups.

| Peptide | Sequence | MW (mg/mmol) | Main function |
| --- | --- | --- | --- |
| RGD | CGGGRGDS | 707,7 | Integrin-binding domain |
| IKVAV | CGGGIKVAV | 802,97 | Laminin-derived |
| YIGSR | CGGGYIGSR | 868,94 | Laminin B1 chain |
| QK | CGGGKLTWQELYQLKYKGI | 2185,51 | VEGF mimetic peptide |

### 2.4 HA characterization

The purified HA-Nb, HA-Nb-Mal and the HA functionalized with peptides were characterized to assess functionalization and purification process.

#### 1H-NMR

The samples were dissolved in phosphate buffered D_2_O (pH 7.4, [phosphate = 56.9 mM, and [3-(trimethylsilyl)-1-propanesulfonic acid-*d*_6_ sodium salt] = 0.11 mM). All ^1^H-NMR spectra were recorded at 299.7 K with a Bruker Avance III HD 700 MHz spectrometer, comprising a cryocooled three-channel TCI probe. The spectra were processed and analyzed with TopSpin 4.0 software (Bruker, Germany). The degree of substitution for Nb and Mal were determined by the ratio between the integral values of the non-anomeric backbone protons in the repeating disaccharide unit of HA and the integral value of either the vinylic Nb group (5.9-6.4 ppm) or the Mal group (6.85-6.95 ppm).

#### Gel permeation chromatography (GPC)

Gel permeation chromatography was used to determine the molecular weight and dispersity of the HA(-derived) polymers before and after the peptide functionalization. Samples were dissolved in a 0.1 M NaNO_3_ solution (1mg/ml), the mobile phase of the system. All samples were filtered through filters with pore size 0.2 µm prior to injecting a volume of 50 µL. Measurements were performed on a Prominence-I LC-2030C3D LC (Shimadzu) system comprising an autosampler, a Shodex SB-G 6B guard (6.0 mm x 50 mm) column, followed by a Shodex SB-803/SB804 HQ (8.0 mm x 300 mm) columns, a refractive index detector, and a photodiode array detector. The flow rate was kept at 1 ml·min^−1^ at T = 25 °C. Calibration was performed using PEG standards up to 545,000 MW (PEG calibration kit, Agilent Technologies).

### 2.5 HA hydrogel formation and characterization

In a typical procedure, HA-peptide_x_ hydrogels were prepared by dissolving the polymer in PBS. Subsequently, the dissolved polymer was mixed with the crosslinker ([-SH] = 25 mM] and the photo-initiator to reach a final polymer concentration of 2% w/v. Lithium phenyl-2,4,6-trimethylbenzoylphosphinate (LAP) was used as photoinitiator [2 mM final concentration]. Two crosslinkers were compared along the study, PEG dithiol (5k Biopharma PEG, HO003003-5K) and a custom-made cysteine terminal MMP2/9 cleavable peptide (GenScript, purity > 96%). The thiol to Nb were reacted in a 1-to-1 equivalent ratio to the Nb groups. Finally, 50 μl of polymer mix was cast in a mold and irradiated with a UV lamp (λ = 365 nm, I = 10 mW/cm^2^, 20s). For the swelling test a silicon mold was used to form the hydrogels. For the cell culture experiments, the gels were formed directly in the wells of a black 96-well plate with transparent glass bottom.

#### Swelling/degradation

All HA gels formulations were tested for swelling over 7 days, by changing the crosslinker. The test was conducted in PBS at 37 °C.

Briefly, HA-peptide_x_ was dissolved in PBS, then the crosslinker and the photoinitiator were added. 50 μl of solutions was cast in a silicon mold and irradiated with a UV light (λ = 365, I = 10 mW/cm^2^, 20s exposure). Hydrogels were produced in triplicates per condition. Subsequently, the samples were placed in pre-weighted containers and submerged with PBS supplemented with PenStrep. At different time points, the solution was discarded and the samples weighed. The water uptake was calculated with the following formula:

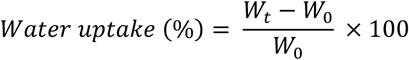

Where W_0_ is the initial mass of the hydrogel immediately after cross-linking and W_t_ is the mass of the gel at each time point.

#### Rheology

Photo-rheology was conducted on all HA-peptide formulations. Gel formulations were prepared according to the earlier described protocol and crosslinked in-situ with PEG/MMP-diSH (n=3). In short, the polymers were dissolved in PBS overnight at 4 °C. Then, the crosslinker and LAP were added to the solution before measuring. The measurements were performed on a DHR2 instrument (TA Instruments), using a cone-plate geometry (20 mm plate diameter, 2.002° cone angle) at 20°C, to mimic crosslinking conditions. The instrument was equipped with a UV light source at 365 nm and the intensity was set at 10 mW/cm^2^. All samples undergone the same 4 step procedure: (1) time sweep for 5 min at 1 rad/s and 1% strain, after the first minute, UV was applied for 1 minute; (2) oscillation frequency 100-0.1 rad/s, 1% strain; (3) oscillation amplitude 1 rad/s, 0.1-1000% strain.

### 2.6 Hybrid construct fabrication and characterization

For the hybrid constructs, melt-extrusion scaffolds were fabricated, comprising medical grade polycaprolactone (PCL, Corbion) and β-tricalcium phosphate (TCP, Kuros Bioscience) in a ratio of 60:40, as described elsewhere [29]. A biomimetic diamond-shape pore geometry was chosen. Scaffolds of 6mm (diameter) x 2mm (height) were printed for biological studies, while 6mm x 4mm samples were used for mechanical testing.

#### Mechanical testing

The mechanical performance of the hybrid constructs upon compression was determined with a TA ElectroForce system (TA Instruments), with a 450N load cell, and controlled with Win7 software. HA-Nb was used to produce hybrid constructs for mechanical testing. The polymer was dissolved in PBS, mixed with PEG/MMP-diSH crosslinker and LAP and cast on the scaffolds placed in a silicon mold. The hydrogels were crosslinked with a UV lamp at 10 mW/cm^2^. The samples were compressed at a strain rate of 0.04 mm/s up to a maximum of 50% compression. Stress values were obtained dividing the measured load against the apparent area of the constructs. The compressive modulus was calculated between approximately 5–15% strain in the linear region of the stress–strain curves. The yield stress and strain values were calculated with the 0.2% offset method. The toughness of the samples was approximated as the area under the curve in the elastic region.

### 2.7 Cell culture

Human mesenchymal stromal cells derived from bone-marrow (hMSCs, PromoCell, lot # 439Z037.1) were expanded at a density of 1,000 cells·cm^−2^ in α-minimum essential medium (α-MEM) (Gibco) supplemented with 10% *v/v* fetal bovine serum (FBS) and ascorbic acid (ASAP). The medium was refreshed every 2-3 days. Cells were passaged at 70-80% confluency and used at passage 5-6 for experiments.

Human umbilical vein endothelial cells (HUVECs, Lonza) were cultured at a density between 6,000-10,000 cells·cm^−2^ in endothelial cell growth medium ready to use kit (Promocell, C-22010). The medium was refreshed every 2-3 days. Cells were used at passage 4-6 for the experiments. HUVEC-GFP (PeloBiotech, PB-CAP-0001GFP) were cultured at a density of 10,000 cells·cm^−2^ in endothelial growth medium (Pelobiotech, PB-CAP-02). The medium was refreshed every 2-3 days. Cells were used at passage 8-9 for the experiments. HUVEC-GFP were used for fluorescent imaging, while normal HUVECs were used for all the other biological assays.

### 2.8 Co-culture conditions validation

hMSCs and hUVECs were co-cultured, at a ratio of both medium and cells of 1:1, based on the literature [37,38]. The culture conditions were validated in 2D, by evaluating the expression of cell specific markers and metabolic activity. The culture conditions are summarized in **Table 2**. Cells were seeded in a treated 48-well plate at a density of 10,000 cells·cm^−2^.

**Table 2.** Cell culture conditions validated in 2D and performed analysis.

| Cell | Medium | Analysis |
| --- | --- | --- |
| hMSCs | BM:EM = 100:0 | Immunostaining: RUNX2, CD31/Ve-Cadherin, DAPI, Phalloidin<br>Metabolic activity: Presto Blue |
| hMSCs | BM:EM = 50:50 |  |
| hUVECs | BM:EM = 0:100 |  |
| hUVECs | BM:EM = 50:50 |  |
| hMSCs:hUVECs | BM:EM = 50:50 |  |

On days 1, 3 and 7, the metabolic activity was measured with a Presto Blue™ (Invitrogen) Cell viability Reagent, according to manufacturer instructions, by incubating the samples in medium at 10% v/v for 15 min. The fluorescence was measured with a plate reader at excitation/emission wavelength 535-615 nm. After the assay, the cells were washed 3 times with PBS and fresh medium was added. Blank medium without cells was used as control.

On day 7, the samples were fixed in 4% v/v paraformaldehyde (PFA) for 10 min and washed in PBS. For the immunostaining, a 10 min permeabilization was first performed with Triton-X 0.1% v/v, followed by a blocking step with BSA 3% w/v and Tween20 0.1% v/v for 1.5 h. Then, primary antibodies and phalloidin were incubated overnight at 4°C. After washing, the secondary antibodies were incubated for 45 min and washed before adding DAPI (1:300) for 20 min. Antibodies and dilutions are reported in **Table S1**. Samples were imaged with a Slide Scanner microscope (Nikon Ti-E microscope, equipped with a Lumencor Spectra light source, an Andor Zyla 5.5 sCMOS camera, a Nikon DS-Ri2 color camera, and an MCL NANO Z200-N TI z-stage).

### 2.9 Cell seeding on top of the hydrogels

#### Cell seeding

The adhesion, viability and metabolic activity of hMSCs and HUVECs were first evaluated by seeding the cells on top of the hydrogels (**Figure S1A**). Four different conditions were tested as shown in **Table 3**. The hydrogels were prepared by casting 50 μl of polymer solution in a well of a black 96-well plate and crosslinked with a UV lamp at 10 mW/cm^2^ for 20 s. Then, the gels were submerged with culture medium supplemented with PenStrep 1% v/v and left to reach swelling equilibrium overnight. Subsequently, the medium was discarded, and the cells were seeded on top of the gels at a density of 1×10^5^ cells·cm^−2^. For each hydrogel, three cell conditions were seeded: hMSCs only, hUVECs only and their co-culture in a ratio of 1:1. HUVEC-GFP were used for imaging experiments.

**Table 3.** Summary of the tested conditions for the seeding on top of the hydrogels.

| HA functionalization | Crosslinker | Tested cells | Cell density |
| --- | --- | --- | --- |
| HA-RGD | PEG | hMSCs | $1 \times 10^5$ cells·cm<sup>-2</sup> |
|  |  | hUVECs |  |
|  |  | hMSCs:hUVECs = 50:50 |  |
| HA-RGD | MMP | hMSCs | $1 \times 10^5$ cells·cm<sup>-2</sup> |
|  |  | hUVECs |  |
|  |  | hMSCs:hUVECs = 50:50 |  |
| HA-YIGSR | PEG | hMSCs | $1 \times 10^5$ cells·cm<sup>-2</sup> |
|  |  | hUVECs |  |
|  |  | hMSCs:hUVECs = 50:50 |  |
| HA-YIGSR | MMP | hMSCs | $1 \times 10^5$ cells·cm<sup>-2</sup> |
|  |  | hUVECs |  |
|  |  | hMSCs:hUVECs = 50:50 |  |

#### Adhesion and viability

Cell adhesion and viability after 24h from the seeding were investigated with brightfield/fluorescent imaging and fixable live/dead immunostaining. Briefly, the samples were washed 2x with PSB, and the Fixable L/D solution was added and incubated in dark for 25 min, according to manufacturer instructions. Then, the solution was washed 2x with PBS and the samples were fixed with 4% v/v PFA for 20 min. The samples were washed again 3X and stored in PBS at 4°C for further staining. Subsequently, DAPI staining was performed. The samples were permeabilized with 0.1% v/v Triton X for 10 min, washed with PBS 3x and stained with DAPI (1/300), before imaging with Slidescanner Microscope. The percentage of alive cells (Live %) was quantified in different conditions, by image analysis with the software *ImageJ*. Briefly, the number of dead and total cells was quantified by counting the dead-positive and Dapi-positive nuclei. Then, the Live% was calculated with the following formula:

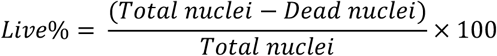

The percentage of alive cells in the RGD and YIGSR conditions were compared. After 24h, the covered area % in all culture conditions was also quantified. The brightfield images were processed on *ImageJ*, to quantify the area occupied by the cells. Then, the cell area was divided by the total area of the image and the covered percentage was calculated. The values were plotted to compare the effect on the peptide on the cell adhesion.

#### Metabolic activity and spreading

On day 1, 3 and 7 the metabolic activity (n=3) was measured by incubating a 10% v/v Presto Blue solution for 15 min, followed by measurements of fluorescence with plate reader. Moreover, on day 7 the samples were also fixed with 4% v/v PFA for fluorescence staining. Fixed samples were permeabilized with 0.1% v/v Triton X for 10 min and phalloidin was incubated overnight at 4°C. Then, the samples were washed with PBS and DAPI staining (1/300) was performed for 20 min. The samples were imaged with a Slidescanner Microscope to investigate cell spreading and morphology after 7 days of culture on different gel conditions. Detailed list of antibodies and dilutions is provided in **Table S1**.

### 2.10 Cell encapsulation in the hybrid scaffolds

#### Cell encapsulation in the hydrogels only

First, the morphology and metabolic activity of cells was evaluated, when encapsulated in HA-RGD or HA-RGD/YIGSR (ratio 1:1) hydrogels only, crosslinked with either PEG or MMP, as summarized in **Table 4**.

**Table 4.** Summary of the tested conditions for cell encapsulation in the hydrogels only and in the hybrid scaffolds.

| HA functionalization | Crosslinker | Tested cells | Cell density |
| --- | --- | --- | --- |
| HA-RGD | PEG | hMSCs | 5·10<sup>6</sup> cells·ml<sup>-1</sup> |
|  |  | hUVECs |  |
|  |  | hMSCs:hUVECs = 50:50 |  |
| HA-RGD | MMP | hMSCs | 5·10<sup>6</sup> cells·ml<sup>-1</sup> |
|  |  | hUVECs |  |
|  |  | hMSCs:hUVECs = 50:50 |  |
| HA-RGD/YIGSR | PEG | hMSCs | 5·10<sup>6</sup> cells·ml<sup>-1</sup> |
|  |  | hUVECs |  |
|  |  | hMSCs:hUVECs = 50:50 |  |
| HA-RGD/YIGSR | MMP | hMSCs | 5·10<sup>6</sup> cells·ml<sup>-1</sup> |
|  |  | hUVECs |  |
|  |  | hMSCs:hUVECs = 50:50 |  |

Hydrogel precursor solutions were prepared and mixed with crosslinkers, LAP and cells, at a density of 5·10^6^ cells·ml^−1^. Then, 50 μl of solution was cast in a black 96-well plate and crosslinked with a UV lamp at 10 mW/cm^2^ for 20s (**Figure S1B**). The gels were submerged with medium, that was replaced every 2-3 days.

On day 1, 3 and 7, a Presto Blue™ assay was performed to evaluate the metabolic activity on the cells over time. Briefly, a 10% v/v of Presto Blue reagent in medium solution was prepared and incubated with the samples for 2h, to allow complete diffusion. Then, 100 μl in duplicates were transferred to a black-bottom 96-well plate and the fluorescent was recorded with a plate reader. The samples were thoroughly washed with PBS and new medium was incubated. Gels without cells were used as blanks, and their fluorescent values were subtracted by the cell signals.

Moreover, on day 1, 3 and 7, cell morphology and organization were monitored with a Live Cell microscope. On day 7, the samples were fixed in 4% v/v PFA and stained with phalloidin and DAPI, as described above. Fluorescent images were taken with a Laser Confocal microscope. List of antibodies and dilutions is reported in **Table S1**.

#### Cell encapsulation in the hybrid constructs

Next, cells were encapsulated in hybrid constructs, comprising a PCL/β-TCP scaffold and HA-RGD or HA-RGD/YIGSR (1:1 ratio) hydrogels, crosslinked with either PEG or MMP (**Table 4**). Prior to cell seeding, the scaffolds were disinfected in EtOH 70% v/v for 20 min, washed 3 times in PBS and incubated overnight in basic cell culture medium (alpha-MEM, 10% v/v fetal bovine serum, 1% v/v ascorbic acid), to favor protein adsorption. Then, the scaffolds were dried on absorbent paper and placed on the bottom of a 96-well plates. Subsequently, hydrogel precursor solutions were mixed with cells (density 5·10^6^ cells·ml^−1^) and 50 μl were injected throughout the scaffolds to reach complete embedding. The hydrogels were crosslinked with a UV lamp (λ = 365 nm, I = 10 mW/cm^2^, 30s) (**Figure S1B**). The constructs were submerged with culture medium that was replaced every 2-3 days.

On day 1,3 and 7, cells morphology was monitored with live imaging on a Live Cell microscope. On day 7, samples were fixed with 4% v/v PFA and a DAPI/phalloidin staining was performed as reported in the previous sections. A Laser Confocal microscope was used to image fluorescent samples. The antibodies and the dilutions that were used are reported in **Table S1**. Fluorescent images of the co-culture conditions were processed with ImageJ software for background removal and subsequently analyzed with the software AngioTool (Java) to quantify vessel formation parameters (i.e. total tubule length, vessel area, and number of junctions). Confocal images from both the top and the cross section of the samples were analyzed. An average of the results from the two views was also calculated. On day 7, qPCR was performed to evaluate gene expression in the hybrid scaffolds.

Hydrogel only samples were used as reference. Briefly, samples were washed with PBS, minced, collected in 1 mL TRIzol (Invitrogen), and stored at −80°C. RNA was extracted with chloroform method, followed by overnight precipitation in isopropanol. RNA concentrations were measured using a BioDrop uLITE instrument (Scientific Laboratory Supplies). cDNA was synthesized using an iScript cDNA Synthesis Kit (Bio-Rad), according to manufacturer’s instructions. Finally, qPCR was performed in 10 μL volume, adding 4 ng of cDNA to the iQ SYBER green Supermix (Bio-Rad) and primer solution. The list of primers is reported in **Table S2**. The 2^−ΔΔCt^ method was used to calculate the relative gene expression.

### 2.11 Statistical analysis

All data were reported as means ± standard deviation. Statistical analysis was performed using GraphPad Prism. Normality of the data was first assessed. Statistical significance was determined by One-way analysis of variance with Tukey’s honest significance test multiple comparison tests. If the data distribution was not normal, a Kruskal-Wallis non-parametric test was performed. (****) p < 0.0001, (***) p < 0.001, (**) p < 0.01, (*) p < 0.05.

## 3. Results & Discussion

### 3.1 Hyaluronic acid-Norbornene-Maleimide synthesis and characterization

HA-Nb-Mal was synthesized via a two-step reaction pathway using DMTMM as activator agent of the carboxylic acid group (**Figure 1A**). The Nb served as addressable unit for PEG-diSH or MMP-diSH to create a hydrogel network via thiol-ene chemistry, whereas the Mal groups were used to graft thiolated peptides via Michael addition reactions (**Figure 1B**).

**Figure 1.**
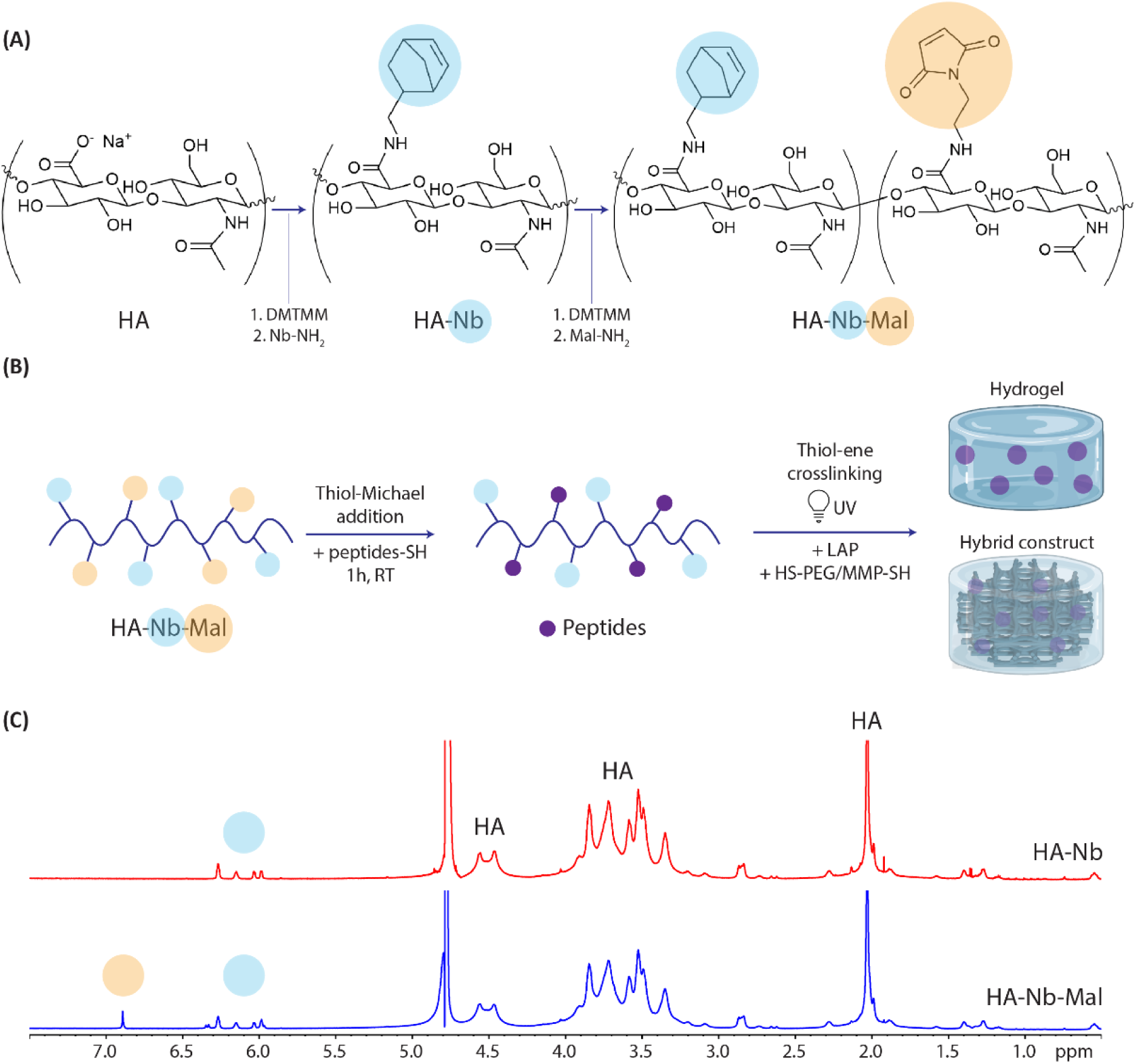
Schematics of reaction steps to obtain the final hydrogels. (A) HA two-steps functionalization where Nb is reacted first, followed by Mal. (B) Covalent grafting of the different peptides to the HA backbone, via Thiol-Michael addition reaction with Mal groups; thiol-ene reaction with UV to crosslink Nb groups and obtain the final functionalized hydrogel. (C) ^1^H-NMR spectra of HA-Nb and HA-Nb-Mal showing the presence of Nb and Mal groups, indicating successful substitution.

The NMR spectrum of the HA-Nb-Mal showed the appearance of the characteristic peak of the double bond of the Nb (5.9-6.4 ppm) and the Mal peak (6.9 ppm). Evaluation of the integral ratio of the peaks compared to the HA backbone revealed a degree of substitution of norbornene of 8±1%, and maleimide 3±1% (**Figure 1C**). Before hydrogel formation, peptides (i.e. RGD, IKVAV, YIGSR, and/or QK) were grafted on the HA-Nb-Mal. The complete disappearance of the Mal peak at 6.9 ppm (**Figure S2A)** as well as a small observed increase in molecular weight compared to pristine HA-Nb-Mal during GPC analysis (**Figure S2B**) suggest successful conjugation of the peptides.

Notably, when the peptides were not pre-grafted before hydrogel formation (see also section 3.2), crosslinking immediately occurred spontaneously. We believe the thiol groups of the crosslinker spontaneously reacted with the Mal groups, prior to UV illumination. The absence of this observation in the pre-grafted system further indicates successful peptide anchoring to the HA backbone.

### 3.2 Hydrogel characterization

After peptide conjugation to the maleimide unit, hydrogel formation was induced via UV using PEG-diSH or MMP-diSH as crosslinker. In preliminary tests, 1% and 2% w/v HA hydrogels were prepared using a 1:1 crosslinker (i.e. the thiol group of the PEG) versus Nb groups (**Figure S3A**). The crosslinking kinetics and viscoelastic properties of the hydrogels were determined by photo-rheology. Compared to the 2% w/v hydrogel, the 1% w/v hydrogel was softer (G’ = 309.5 ± 123.1 Pa vs 1699.0 ± 315.9 Pa), did not maintain a good shape (i.e. was difficult to handle), and displayed fast degradation (< 7 days) when immersed in PBS. Thus, 2% w/v hydrogel formulation was used in the remainder of the study.

Next, the properties of HA functionalized with different peptides (i.e. HA-RGD, HA-IKVAV, HA-YIGSR, HA-QK) hydrogels and crosslinked with two different crosslinkers (i.e. PEG-diSH and MMP-diSH) were compared. Prior to applying UV, all gels remained a liquid except the HA-QK formulation that immediately gelated as indicated by the value of G’ being one order of magnitude above G” (**Figure 2A-B**). We attributed this to the significantly larger size of the peptide, resulting in a higher entanglement of the network, or electrostatic interactions resulting from the presence of charged amino acids. In addition, we observed insoluble filaments in the hydrogels after crosslinking. Regardless, uncontrolled spontaneous gelation prevented adequate handling of the formulations with QK, and these were excluded from further biological studies.

**Figure 2.**
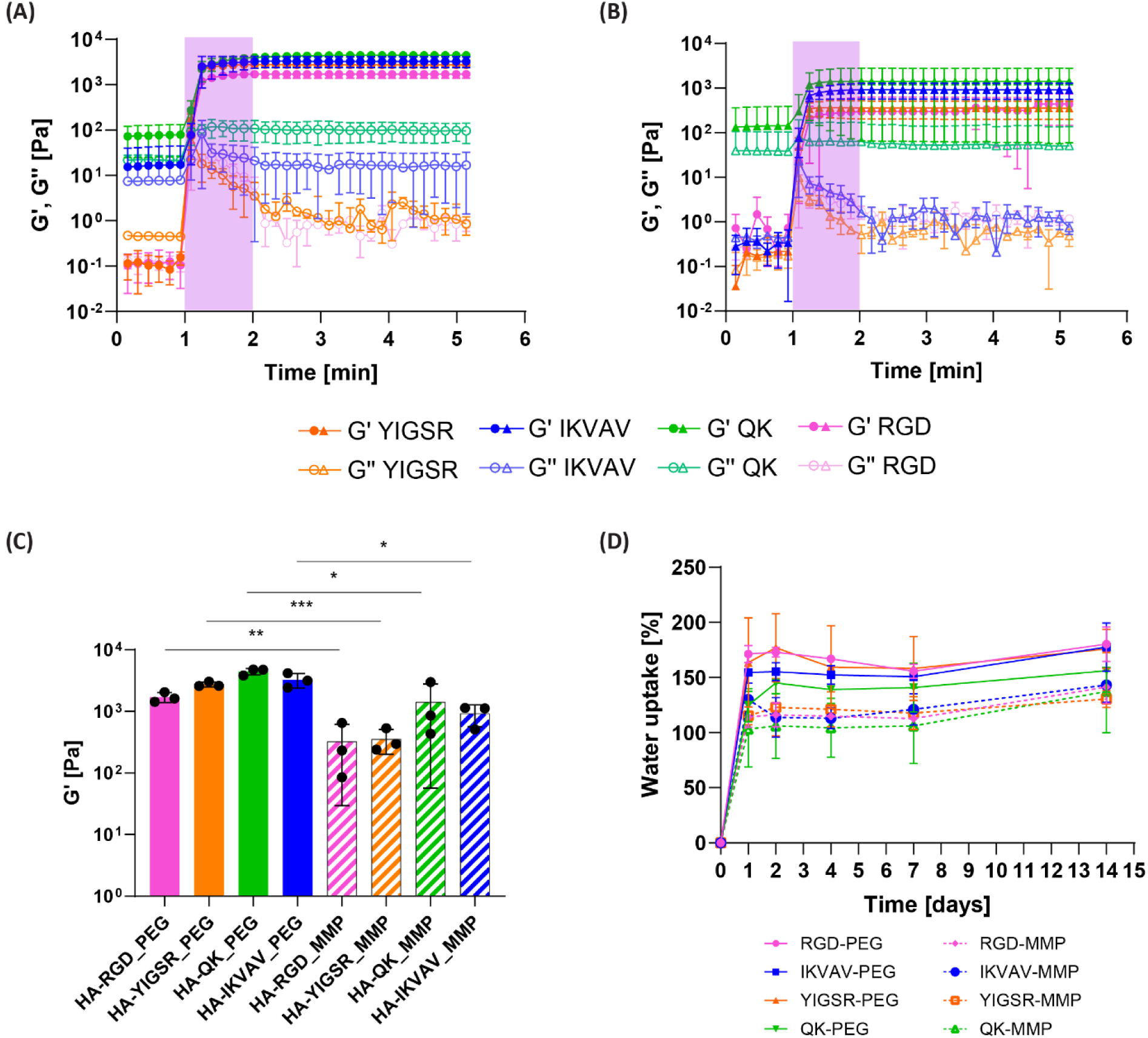
Characterization of the hydrogels. (A) Time sweep step of HA-peptide-PEG and (B) HA-peptide-MMP, showing storage and loss moduli, and the time window where the UV light was applied. (C) Storage moduli values of different hydrogel formulations showing the effect of peptide and crosslinker variations. (D) Water uptake percentage of different hydrogel formulations, varying the peptide and the crosslinker. Statistical significance represented by: ***<0.001, **<0.01, *<0.05.

When UV irradiation was applied to all formulations, G’ values rapidly increased, reaching a plateau where G’ > G’’ in less than 20 seconds (**Figure 2A-B**). This observation implies that the coupled peptides did not interfere with the thiol-ene crosslinking reaction. The stiffness of the hydrogels was extrapolated from the plateau (**Figure 2C**). The MMP-crosslinked groups displayed significantly lower G’ values compared to their respective PEG-crosslinked group (e.g. HA-RGD-PEG G’ = 1699.0 ± 315.9 Pa vs HA-RGD-MMP G’ = 321.8 ± 292.5 Pa). Since the MMP-diSH is much shorter than the PEG-diSH (1041.25 g/mol vs 5000 g/mol), we hypothesized that intramolecular reactions occurred more in the MMP conditions, leading to less active crosslinkers that contribute to the stiffness of the network. When comparing the formulations comprising the same crosslinker, no significant differences were detected and the stiffness ranged from 1699 ± 315.9 to 4440 ± 527.8 Pa for the PEG group and from 321.8 ± 292.5 to 1423 ± 1366 Pa for the MMP group. Overall, all the values were in the range 10^2^ - 10^3^ Pa, comparable with most soft tissues in the body [39–41].

The frequency sweep results showed that all formulations displayed frequency-independent moduli over the measured range (1-100 rad/s) (**Figure S3B-C**). In addition, the strain sweep showed typical elastic breaking behavior, with all hydrogel networks breaking around 200% (**Figure S3D-E**). This elastic behavior is typically observed in covalently crosslinked networks.

The swelling behavior of different hydrogel formulations was tested in physiological conditions by incubating them in PBS at 37°C and refreshing the PBS every 2-3 days to simulate cell culture. All gels swelled up to around 150-200% within 24h from incubation (**Figure 2D**). While no effect was reported by varying the grafted peptide, a lower swelling was observed for the MMP group compared to the PEG one (average of 158.9 ± 12.73% versus 119.3 ± 5.51%) (**Figure S3F**). The shorter length of the MMP crosslinker might result in smaller mesh size of the network, allowing lower water uptake [42]. This result aligns with what observed for the rheological behavior.

### 3.3 Hybrid construct characterization

Hybrid constructs comprising of the PCL/β-TCP scaffold and a HA-peptide_x_-crosslinker_y_ were fabricated via the injection of a liquid formulation throughout the AM construct, followed by UV crosslinking for 30 sec to ensure complete crosslinking. The injection volume was optimized to 50 µL to ensure complete filling of the pores through all layers of the scaffold, as shown by the stereomicroscope images of both the top view (**Figure 3A, Figure S4A**) and the cross-section (**Figure 3B**).

**Figure 3.**
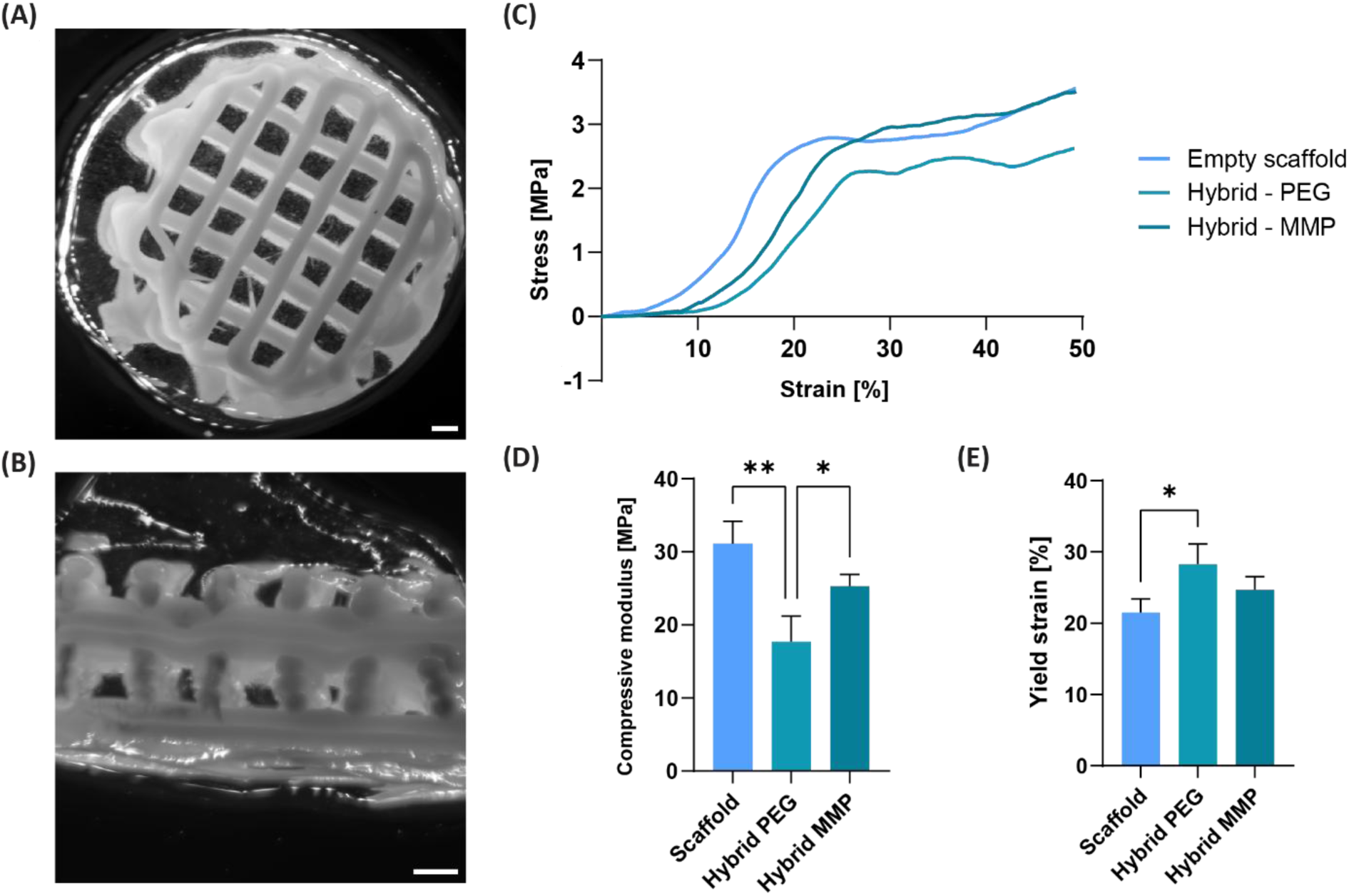
Characterization of hybrid constructs. Stereomicroscope image of the hybrid constructs, comprising the scaffold and the HA hydrogel, (A) from the top and (B) from the bottom. Scale bar = 500 μm. (C) Representative stress-strain curves of the scaffold only, and hybrid scaffolds crosslinked with either PEG or MMP. (D) Compressive moduli [MPa] and (E) Yield strain [%] values comparing the different conditions. Statistical significance represented by: **<0.01, *<0.05.

Since no major differences in elastic properties of hydrogels were observed during rheology among the different peptide groups, we subjected only HA-Nb gels formulation (PEG or MMP) to a compression test. The mechanical properties of the hybrid constructs were compared to the empty scaffolds. The stress-strain curves showed a similar behavior for all conditions, with the scaffold being dominant in the mechanical behavior. This observation is well-reported in the literature, where hydrogels are reinforced to enhance their overall mechanical behavior of several magnitudes [43,44]. All curves presented an initial elastic response, followed by a plastic deformation during the plateau phase, and finally a second modulus increase due to pore closure (**Figure 3C**). The extended toe region of the hybrid construct is likely a result of the slight overfilling of the hydrogels. The following compressive moduli were determined for the constructs: E_empty scaffold_ = 31.1 ± 3.1 MPa, E_Hybrid-PEG_ = 17.7 ± 3.5 MPa, E_Hybrid-MMP_ = 25.3 ± 1.6 MPa (**Figure 3D**). Although they are within the same regime, the addition of the hydrogel slightly reduced the compressive moduli of the construct. This observation could be explained by the dissipation of load from the scaffold to the soft gel. Also, the hydrogel may have introduced a dampening effect in the hybrid constructs, further indicated by a shift of the curves towards higher yield strain values (**Figure 3E**). Toughness and yield stress did not display significant differences (**Figure S4B-C**).

### 3.4 Cell co-culture validation

hMSCs and hUVECs have often been co-cultured to promote the formation of mature and vascular network. These cells interact through paracrine signaling of growth factors, such as VEGF and IGF, and through gap junctions, leading to the spontaneous formation of vessels [45]. Based on previous literature studies, we selected a 1:1 cell and culture medium ratio for the co-culture of hMSCs and hUVECs [45–47]. These parameters were first verified in 2D by seeding hMSCs alone, hUVECs alone, or both cells in a 1:1 cell ratio. Subsequently, their characteristics markers, i.e. RUNX2 and CD31/VE-Cad, were evaluated through immunostaining and the metabolic activity was measured (**Figure S5**).

hMSCs and hUVECs expressed their specific markers, both in monoculture and co-culture (**Figure S5A-E**). All the culture conditions displayed an increasing metabolic activity over time; hMSCs appeared more metabolically active at all timepoints than hUVECs (**Figure S5F**). In conclusion, the selected cell and medium ratio (1:1) were shown to permit cell proliferation and expression of specific markers. These culture conditions were used for all further biological characterization.

### 3.5 Cell adhesion and metabolic activity on top of the gels

An initial biological study was conducted to evaluate the ability of HA hydrogels to support adhesion, proliferation, and metabolic activity of hMSCs, HUVECs, and a co-culture thereof. RGD and YIGSR hydrogels were selected for this experiment, crosslinked with either PEG or MMP, as summarized in **Table 3**. RGD is known for its adhesive properties for cells, while YIGSR – a laminin-derived peptide – has been shown to promote tubule formation in hUVECs [48,49]. IKVAV, despite also being a laminin-derived peptide, showed less stable tubule networks and lower total tubule length both *in vitro* and *in vivo*, and was therefore excluded from further studies [50]. Similarly, QK, a VEGF-mimetic peptide, promoted lower tubule formulation when hUVECs were seeded on Matrigel, compared to YIGSR and RGD hydrogels [51]. This, in combination with the practical handling problems reported in the rheological section, led to the exclusion of this peptide from further biological studies.

Cells were seeded on top of the different hydrogel formulations. After 1 day, cells on the RGD condition displayed good adhesion and adopted an elongated morphology for both the co-culture (**Figure 4A**) and the mono-cultures (**Figure S6A**). The covered area % of cells in different conditions showed greater coverage in the RGD gels, both PEG (RGD 12-24% vs YIGSR 5-9%) (**Figure 4B**) and MMP (RGD 18-21% vs YIGSR 6-10%) (**Figure 4C**). This data suggests that cells spread less on the gels comprising YIGSR.

**Figure 4.**
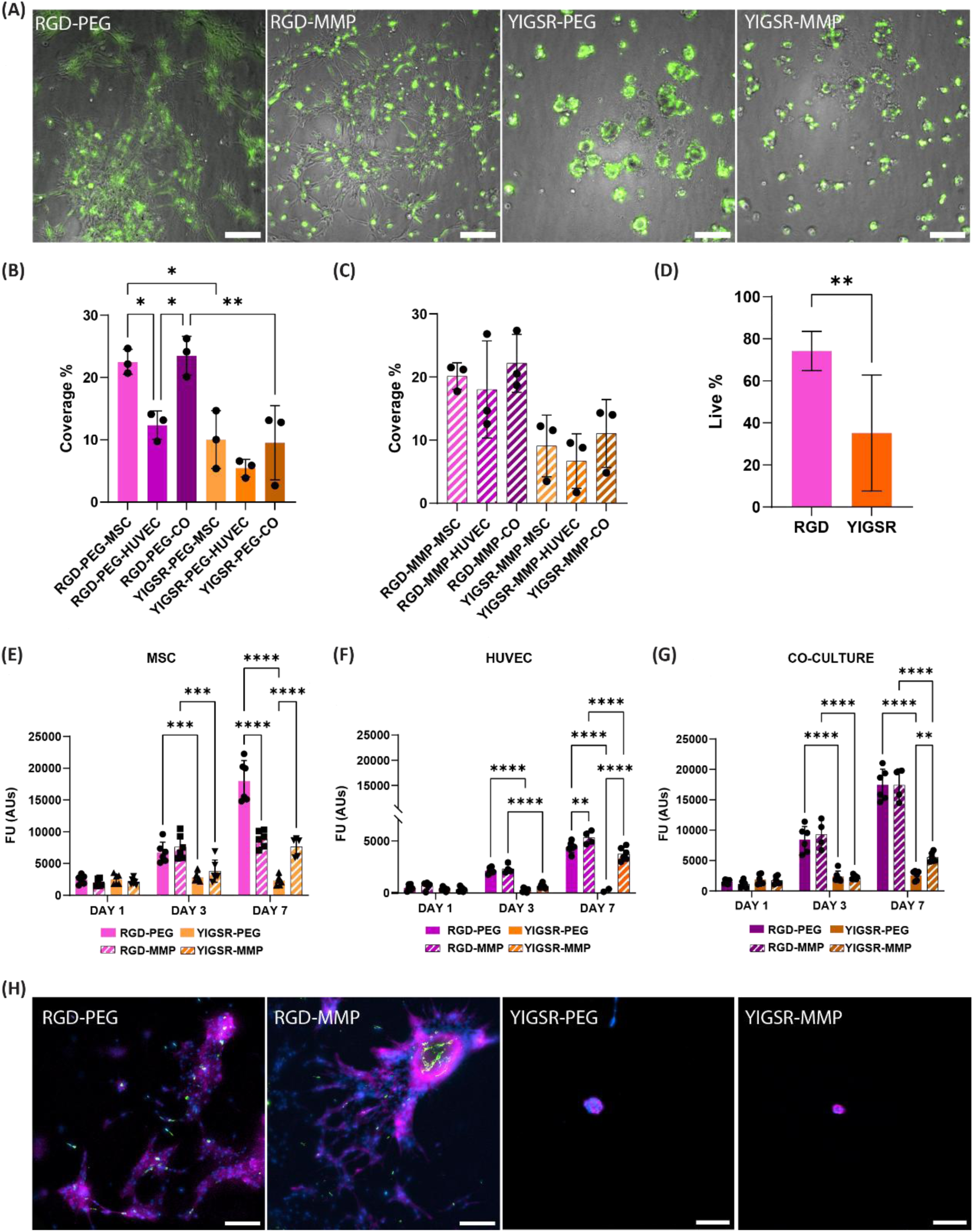
Assessment of cell response when seeded on top of HA-RGD-PEG/MMP or HA-YIGSR-PEG/MMP hydrogels. (A) Qualitative evaluation of the morphology of the co-culture (hMSCs=brightfield, hUVEC-GFP=green) after 24h from the seeding on top of the gels, in different conditions. Scale bar 200 μm. Coverage% of the cells seeded on top of the gels in (B) PEG and (C) MMP condition. (D) Quantification of the live cell% from the Dapi/dead staining, comparing RGD vs YIGSR group, to see the effect of the peptide. Metabolic activity over 7 days of (E) hMSCs, (F) hUVECs and (G) their co-culture, when seeded on different gel conditions. (H) Immunostaining of the co-culture after 7 days on different gel conditions, showing spheroid-like structures on the YIGSR conditions. DAPI (blue), Phalloidin (magenta), hUVEC-GFP (green). Scale bar 200 μm. Statistical significance represented by: ****<0.0001, ***<0.001, **<0.01, *<0.05.

In addition, the quantification of the Live cell % displayed a significantly greater number of alive cells in the RGD gels compared to the YIGSR conditions (74.22 ± 3.80% vs 52.83 ± 2.54%) (**Figure 4D**). This is likely associated with the higher adhesive properties of the gel comprising RGD and not due to toxicity of the gel itself. In YIGSR conditions, the spheroid-like structures of the co-culture comprised a necrotic core. When only hUVECs were seeded on the gels with YIGSR, no cells were left after the staining procedure. These results indicate that the hUVECs do not bind to the YIGSR moiety, preferring RGD instead (**Figure S6B**). No differences were appreciated between the PEG and MMP crosslinkers.

After 7 days, the cells displayed increasing metabolic activity, particularly for the RGD gels, and with around 40-fold higher values for the hMSCs compared to hUVECs in the RGD-PEG condition (**Figure 4E-G**). The higher amount of cells in the microscopy images revealed that the RGD gels supported proliferation. The cells on YIGSR gels maintained their spheroid-like shape, further confirming the preference for RGD (**Figure 4H, Figure S6C**).

The results of this first biological study indicated that the compositions comprising RGD peptides supported better cell adhesion, confirming the central role of the RGD complex in the integrin-mediated cell attachment [52]. On the contrary, the YIGSR hydrogels resulted in lower spreading and cells displaying spheroid-like morphology. Moreover, for the same cell condition, no significant differences were observed when comparing the two crosslinkers (e.g. RGD-PEG-MSC vs RGD-MMP-MSC) (**Figure S7**). Thus, RGD-PEG/MMP conditions were moved to further studies, while the YIGSR hydrogels were excluded. Additionally, a second hydrogel formulation was introduced, HA-RGD/YIGSR crosslinked with either PEG or MMP, comprising a combination of the two peptides in a 1:1 ratio. Here, the synergistic effect between the adhesive properties of RGD and the angiogenic properties of YIGSR was investigated [50].

### 3.6 Cell encapsulation in the hydrogels

After a preliminary cell seeding on top of the gels, we encapsulated the cells within the selected hydrogel formulation (i.e. HA-RGD and HA-RGD/YIGSR, crosslinked with either PEG or MMP) to evaluate their behavior in 3D. Three different cell configurations (**Table 4**) were encapsulated by mixing the cells with the hydrogel precursor solutions, followed by casting and crosslinking with a UV lamp. Cell morphology was monitored using GFP-positive hUVECs and metabolic activity was measured over 7 days. All hydrogel formulations enhanced the metabolic activity of the cells over time. For all cell types, RGD hydrogels promoted significantly higher metabolic activity than RGD/YIGSR ones. While comparing the crosslinkers, MMP performed better than PEG (**Figure 5A, Figure S8A-B**). On day 7, cells adopted an elongated morphology in all hydrogel conditions, suggesting attachment to the hydrogels (**Figure 5B**). When comparing the behavior of hUVECs(-GFP) versus hMSCs in the co-culture conditions, differences were observed: hUVECs remained isolated, whereas hMSCs formed endothelial-like networks throughout the gels.

**Figure 5.**
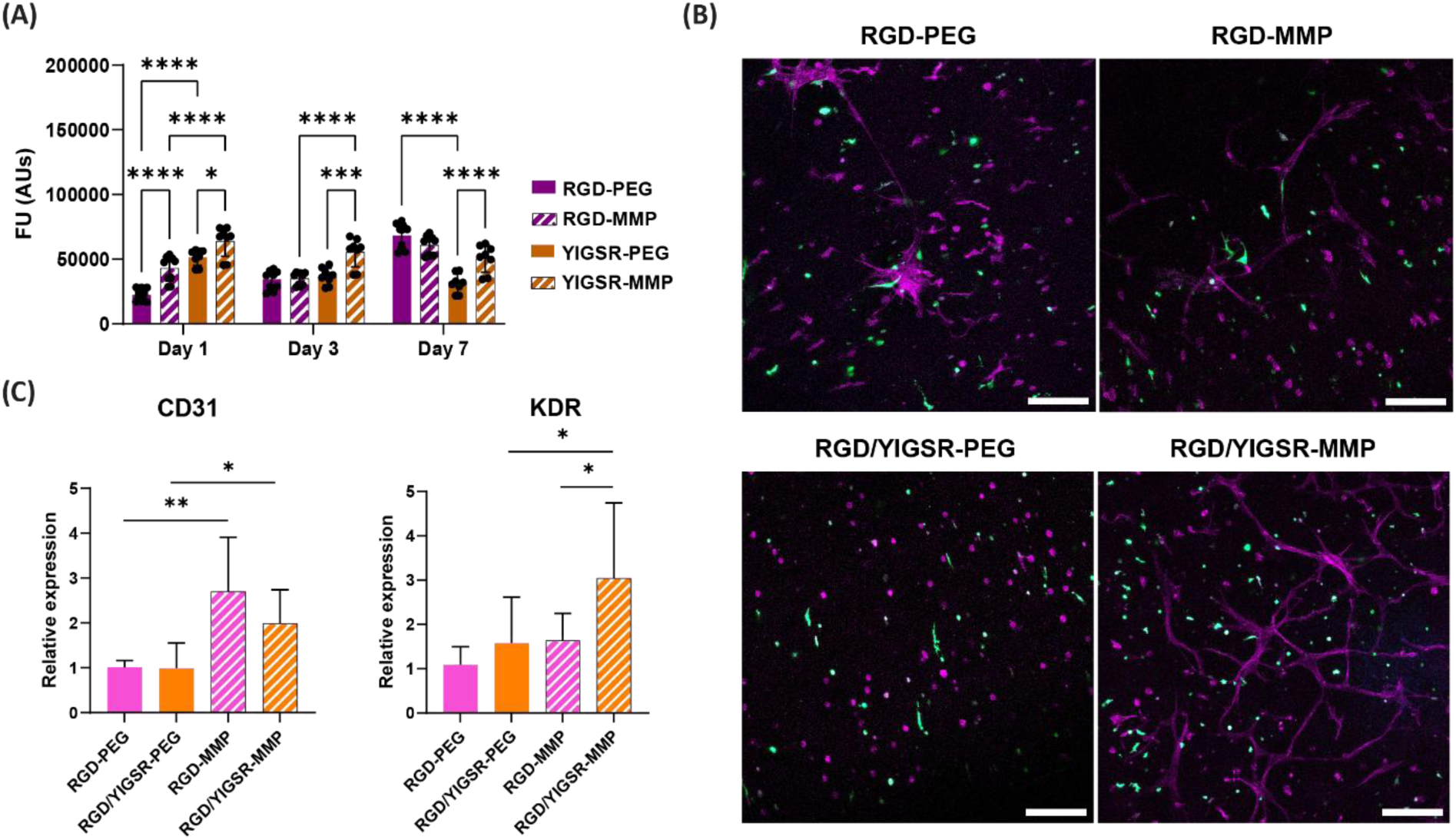
Assessment of the cell response when encapsulated in HA-RGD-PEG or MMP or HA-RGD/YIGSR-PEG or MMP hydrogels. (A) Metabolic activity over 7 days of hMSCs and hUVECs co-culture. (B) Immunostaining of the co-culture conditions after 7 days of culture in different gel conditions. DAPI (blue), Phalloidin (magenta), hUVEC-GFP (green). Scale bar 200 μm. (C) Gene expression of CD31 and KDR genes in the hydrogels only for the co-culture condition. Statistical significance represented by: ****<0.0001, ***<0.001, **<0.01, *<0.05.

Similar behavior was reported before, where hMSCs were shown to undergo mesenchymal-to-endothelial transition when exposed to endothelial cell culture medium in an ECM-like environment [53–55]. For instance, hMSCs encapsulated in a Dextran-HA hydrogel loaded with VEGF or in PEG-based hydrogel showed upregulation of CD31 and KDR after 21 days of culture, and significant matrix remodeling capacity. In the same studies, hUVECs failed to sprout in the hydrogel and presented reduce metabolic activity [56,57]. More recently, Perin et al. reported that hMSCs encapsulated in alginate-based hydrogels also formed extensive endothelial-like network, while hUVECs remained isolated in the hydrogel [30]. Indeed, when we performed gene expression analysis of cells encapsulated in different hydrogel combinations, upregulation of CD31 and KDR was observed in all cell conditions (**Figure 5C, Figure S8C**).

### 3.7 Cell encapsulation in the hybrid constructs

#### 3.7.1 Vascularization

Next, the hydrogel precursors were mixed with cells, and dispersed throughout the scaffold, according to the steps earlier described. After 7 days, vessel formation in the hybrid constructs was evaluated via fluorescence microscopy. To enable qualitative visualization of endothelial and stromal cells, hUVEC-GFP cells were used in the experiment, as previously reported in similar hydrogel-based culture models [58]. Constructs with only hMSCs or only (GFP-)hUVECs did not display any vessel formation. hMSCs appeared to migrate towards the printed fibers while degrading the hydrogels, whereas hUVECs remained isolated in the hydrogel maintaining a rounded morphology. The failure of endothelial cell monocultures to assemble into capillary-like structures reinforces the importance of cellular crosstalk with stromal cells [59]. The combination of paracrine signaling and cell-to-cell interaction leads to recruitment of stromal cells, which are involved in matrix remodeling and stabilization of the newly formed vascular network [10].

Further supporting the hypothesis, in our study the hybrid constructs with all gel formulation comprising a co-culture induced endothelial tubule-like cell organization. Notably, from both a qualitative and quantitative assessment of fluorescent images, the RGD-PEG conditions supported the development of significantly longer capillary-like structures throughout the scaffold thickness (**Figure 6A, Figure S10**). We observed that hMSCs were located both in proximity of the newly formed endothelial structure, supporting the early formation of tubules, and on scaffold fibers. In addition, RGD-PEG scaffolds induced significant upregulation of αSMA gene in the co-culture condition compared to the other scaffolds (**Figure 6B**). αSMA gene expression was used as a preliminary indication of pericyte-like behavior of hMSCs during the early phases of co-culture organization. This phenotype enables hMSCs to support and stabilize vascular structures, by organizing around them [60]. This was shown to keep the network stable up to 130 days *in vivo* [61]. Similar results were reported when a co-culture of hUVECs and hMSCs was seeded on poly(LLA-co-DXO) scaffolds, where αSMA positive hMSCs were identified in proximity to the newly formed tubules [62]. Hence, the upregulation of αSMA gene together with qualitative assessment of cellular organization in the RGD-PEG hybrid scaffolds was considered an initial indication of commitment of hMSCs towards pericyte-like behavior and support the observation of early vessel formation.

**Figure 6.**
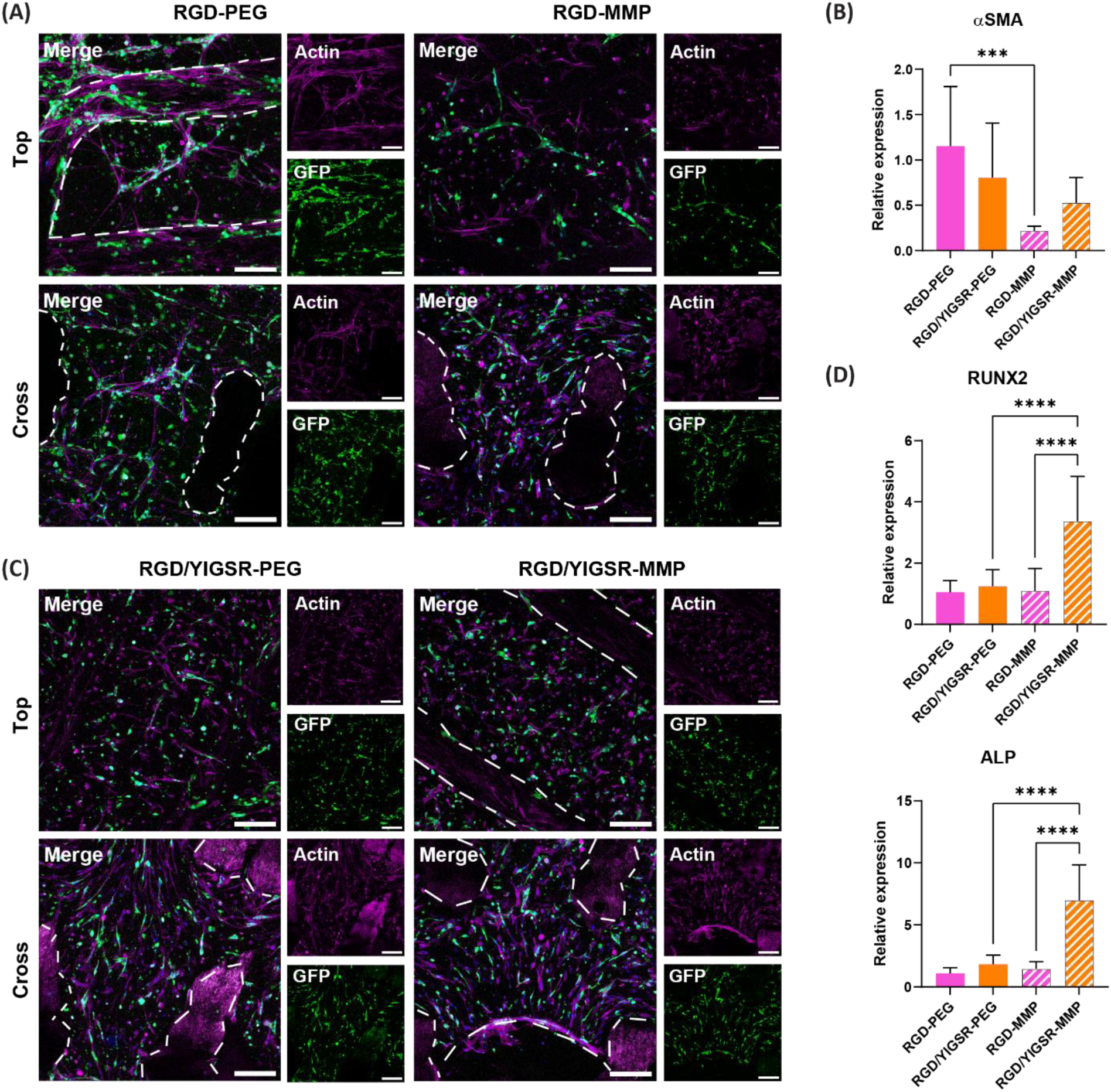
Assessment of cellular response in the hybrid constructs. (A) Fluorescent images (hUVEC-GFP=green, Phalloidin=magenta) of the co-culture condition in RGD-PEG or RGD-MMP hybrid scaffolds. Scale bar 200 μm. (B) Relative gene expression of αSMA in the hybrid constructs with co-culture. Values normalized against the RGD-PEG condition. (C) Fluorescent images (hUVEC-GFP=green, Phalloidin=magenta) of the co-culture condition in RGD/YIGSR-PEG or RGD/YIGSR-MMP hybrid scaffolds. Scale bar 200 μm. (D) Relative gene expression of RUNX2 and ALP in the hybrid constructs with co-culture. Values normalized against the RGD-PEG condition. White striped lines indicate the fibers of the scaffolds. Statistical significance represented by: ****<0.0001, ***<0.001.

Zooming in on the RGD/YIGSR-hybrid constructs, the capillary-like structures appeared shorter and less developed than in the respective RGD scaffolds (**Figure 6C, Figure S10**). Similarly, the hMSCs appeared less spread and only limited contact with endothelial cells was observed. When comparing the two crosslinking methods, significantly lower tubule length, vessel area and number of junctions were recorded only for RGD-MMP samples, which could be attributed to differences in hydrogel degradability that might have resulted in a less stable matrix for cellular anchorage.

In this study, we focused on capturing the initial phases of cellular spatial organization within the hybrid system over a 7-day culture period. For this reason, αSMA gene expression was used as an early indicator of pericyte-like behavior of hMSCs, allowing us to evaluate the effect of the different hydrogel chemistries on the expression of pericyte-related markers. In future studies, the expression of additional markers should be evaluated to further confirm a fully developed pericytic phenotype, including the presence of contractile markers such as PDGF-β or NG2, as well as the formation of a lumen and CD31/αSMA co-localization [28,63,64]. However, these markers are typically observed at later stages of vascular network maturation, which were not reached within the scope of this study. Future studies could therefore aim to assess longer culture periods to evaluate full network functionality.

#### 3.7.2 Osteogenic capacity

Next to promoting vascularization, hybrid scaffolds were designed to possess osteogenic properties, to simultaneously support bone formation. To this end, we have included in the constructs an AM scaffold comprising PCL/β-TCP, which has been shown to promote osteogenesis in earlier studies [29]. Gene expression of early osteogenic markers (i.e. RUNX2 and ALP) was assessed. Both genes were upregulated in thy hybrid constructs when compared to the hydrogels alone, supporting the hypothesis that the scaffold embedded in the gel matrix, not only contributes to mechanical reinforcement, but also guides cells (i.e. hMSCs) towards an osteogenic lineage (**Figure S9A**).

When comparing the gene expression across hybrid scaffold conditions, we observed a 4-fold upregulation of RUNX2 and 6-fold upregulation of ALP in the RGD/YIGSR-MMP hybrid scaffolds (**Figure 6D**). The cells in the respective RGD/YIGSR-PEG only presented around 2-fold upregulation of osteogenic genes, suggesting that the strongest osteogenic properties might arise from the combination of the scaffold and the degradable MMP crosslinker in the hydrogel. These observations highlight that the hydrogel composition is relevant and can be tailored to guide vascularization and osteogenic processes.

Our findings align with other literature studies. For instance, Sousa et al. showed that PCL-nHA scaffolds embedded in alginate/collagen-based hydrogels improved mineralization of hDPSCs spheroids *in vitro* and significant tubule formation *in vivo* when implanted in a subcutaneous rat model [65]. Dubey et al. fabricated melt-electrowritten PCL scaffolds reinforced by a GelMA-based hydrogel and demonstrated that the direct co-culture of hMSCs and hUVECs significantly improved mineralization and upregulated the expression of several osteogenic genes [66]. We hypothesize that the presence of stiffer AM scaffolds with defined fiber orientation provides physical cues that in combination with cellular crosstalk and a defined hydrogel chemistry promote osteogenic differentiation.

Taken together, our results highlight that embedding AM scaffolds within hydrogels with different peptides and crosslinkers influences both endothelial organization and early osteogenic gene expression. Although no unique formulation seemed to simultaneously maximize both outcomes, the results provided insights into how the system can be tuned to guide specific cellular behavior. These effects can be attributed to the ECM-like component provided by the hydrogel, comprising chemical cues (e.g. peptides or crosslinkers) that can guide cell adhesion and matrix remodeling. Moreover, the composite scaffold component provides mechanical and spatial cues to guide cells towards osteogenic differentiation [28,67,68]. However, differences in cell response were observed when comparing the different hydrogel chemical compositions. First, RGD constructs seemed to promote earlier and longer tubule formation, than the respective RGD/YIGSR scaffolds. We hypothesized that the higher RGD concentration in the RGD conditions (double than the mixed peptide) led to faster and greater cell adhesion to the hydrogel and subsequent matrix remodeling. Indeed, other studies demonstrated that cell adhesion to hydrogel matrices is highly dependent on RGD peptide concentration [69]. Instead, the RGD/YIGSR formulations appeared to be more involved in osteogenic commitment, particularly the MMP condition. This could result from a combined effect of lower concentration of adhesive peptide and matrix degradability. In fact, the MMP crosslinker is highly degradable and this could lead to greater migration of cells towards the scaffold fibers.

In the context of pre-vascularization, Matrigel^®^ or GelMA have often been identified as benchmark hydrogels due to their inherent bioactivity and their well-documented ability to support vessel formation [70,71]. However, both present limitations that complicate systematic investigating of individual biochemical cues (e.g. peptides or crosslinkers), as well as their translational potential. Matrigel^®^, being animal-derived, presents high batch-to-batch variability and an undefined biochemical composition [72]. GelMA, despite being more controlled and characterized, lacks modularity, limiting for example the ability to independently control peptide functionalization or crosslinking chemistry. Therefore, the complexity of the elements that can be incorporated is limited. In contrast, the HA-based hydrogel developed in this study was specifically designed for precise control and tunability of matrix properties through peptide coupling or crosslinking. This provides a more defined and systematic platform to study cellular response to the different biochemical and mechanical cues.

Next to the importance of the matrix properties, our results also highlighted the importance of cellular crosstalk in guiding the tubule-like organization. While the hybrid scaffolds encapsulating monocultures of either hMSCs or hUVECs did not display any vessel formation, their co-culture instead promoted early cellular organization. These preliminary findings support the involvement of paracrine interactions between the two cell types. However, a deeper investigation of key pathways (e.g. VEGF, PDGF-β, PI3K/AKT) and their downstream factors could be performed in future studies, for instance through secretome analysis, to elucidate the molecular mechanisms involved in cell-cell communication within the hybrid scaffolds.

Ultimately, the goal is to implant a pre-vascularized construct *in vivo* that can successfully support vessel inosculation and maturation and bone regeneration. These constructs could be used as alternative solution to autologous fibula flap grafts or as a complementary approach in large critical-sized defects. Specifically, pre-vascularized hybrid systems could be applied to enhance vascular coupling and tissue remodeling at the graft-host interface, where rapid inosculation and bone healing are critical for long-term success. In this study, we focused on discovering the pre-vascularization and osteogenic capacity of a novel hybrid construct from a more fundamental angle. Future studies should investigate complete cellular maturation and vascular network stability by performing longer cultures (e.g. 28 days), where expression of late osteogenic markers (e.g. Collagen I, Osteocalcin) and vascular maturation markers (e.g. Collagen IV, αSMA) could be assessed [73–75]. In addition, future research could focus on evaluating different cell seeding regimens, where endothelial and stromal cells are seeded sequentially, which was shown in the literature to be more effective in the maturation of the vascular network. This would better resemble the *in vivo* tissue repair dynamics, where first endothelial cells are activated and then, through the release of growth factors, stromal supportive cells are recruited [76,77].

## 4. Conclusion

In conclusion, this study demonstrated that hybrid scaffolds comprising an AM osteogenic scaffold and a peptide-functionalized HA hydrogel were able to support *in vitro* early vascular-like network formation and osteogenic commitment, when co-culturing endothelial and stromal cells. We successfully demonstrated that the biological outcome of the system can be controlled by tuning the chemistry of the hydrogel, in terms of grafted peptides and crosslinking sequence. Future studies will aim at validating these findings in longer cell culture studies and *in vivo* subcutaneous implantation in a mouse animal model to assess network maturation and inosculation capacity.

## Acknowledgements

The authors would also like to acknowledge the European Research Council starting grant “Cell Hybridge” for financial support under the Horizon2020 framework program (Grant #637308). This project has also received funding from the European Union’s Horizon 2020 FET Open programme under grant agreement No. 964452 (BIRDIE project).

