## Supplementary Information for "Engineering a hybrid 3D construct for bone regeneration to promote simultaneous pre-vascularization and osteogenic differentiation in vitro"

**Table S1.** List of antibodies and dilutions that were used in 2D and 3D experiments.

| Antibody | Species & brand | Dilution |
|---|---|---|
| **2D experiment** | | |
| Phalloidin | Alexa Fluor 488 - Thermofisher | 1:500 |
| Runx2 | Rabbit – Abcam ab192256 | 1:300 |
| CD31 | Mouse – Abcam ab24590 | 1:400 |
| Ve-Cad | Mouse – R&D systems af938 | 1:400 |
| **Seeding on top of the gels** | | |
| Phalloidin | Alexa Fluor 647 - Thermofisher | 1:500 |
| **Encapsulation in the gels/hybrid constructs** | | |
| Phalloidin | Alexa Fluor 647 - Thermofisher | 1:500 |

**Table S2.** Primer sequences of the genes, used for qPCR experiment: runt-related transcription factor (RUNX2), alkaline phosphatase (ALP), platelet endothelial cell adhesion molecule 1 (PECAM-1), kinase insert domain receptor (KDR), actin alpha 2 (ACTA2), beta-2-microglobulin (B2M, housekeeping gene).

| Gene | Forward primer 5' to 3' | Reverse primer 5' to 3' |
|---|---|---|
| RUNX2 | TCAACGATCTGAGATTTGTGGG | GGGGAGGATTTGTGAAGACGG |
| ALP | ACAAGCACTCCCACTTCATC | TTCAGCTCGTACTGCATGTC |
| PECAM1 (CD31) | GAAAGCCAAGGCCAAGCAGATG | TTTCCACGGCATCAGGGACAG |
| KDR (VEGFR2) | CCCTACAAGACCAAAGGGGCAC | GCGATGCCAAGAACTCCATGC |
| ACTA2 ($\alpha$SMA) | ACGTGGGTGACGAAGCACAG | GGGCAACACGAAGCTCATTGTA |
| B2M | ACAAAGTCACATGGTTCACA | GACTTGTCTTTCAGCAAGGA |

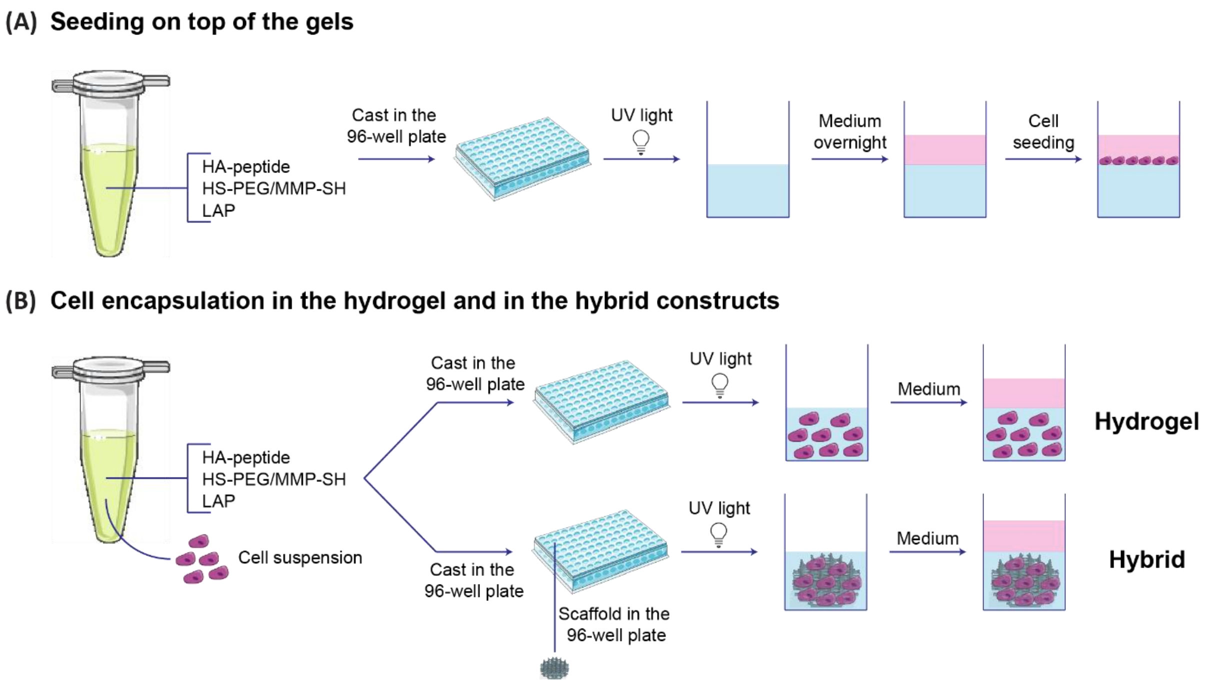


***Figure S1.*** *Schematics of the biological experiments conducted to validate the hydrogel and hybrid constructs. (A) Steps for the cell seeding on top of the gels. (B) Steps for the cell encapsulation in the hydrogel or in the hybrid constructs.*

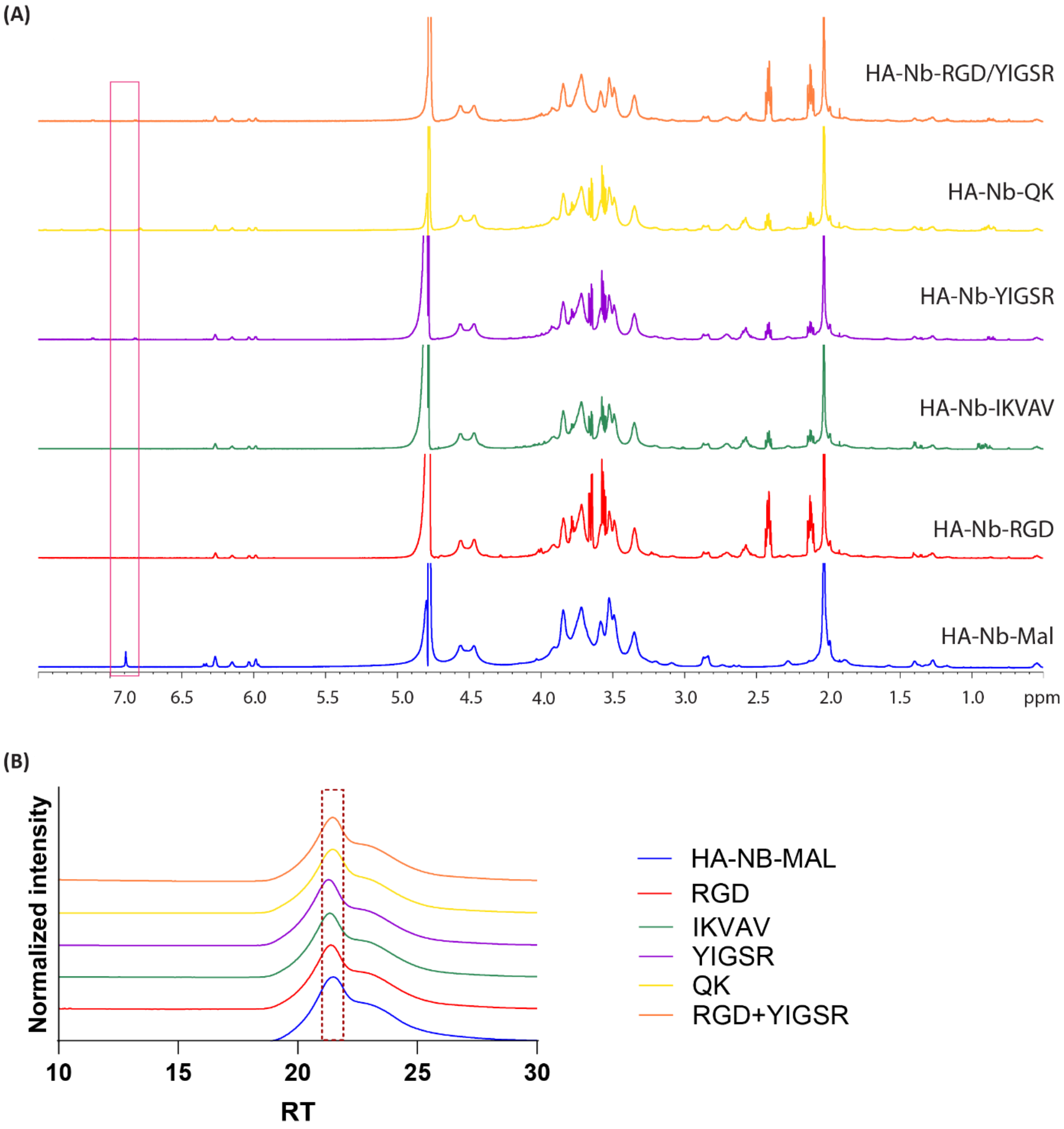


***Figure S2.*** *Characterization of HA after peptide functionalization. (A) NMR of the polymer after peptide grafting, showing that the Mal disappears. (B) GPC of the polymer with and without peptides, showing slight molecular weight increase when the peptides are grafted.*

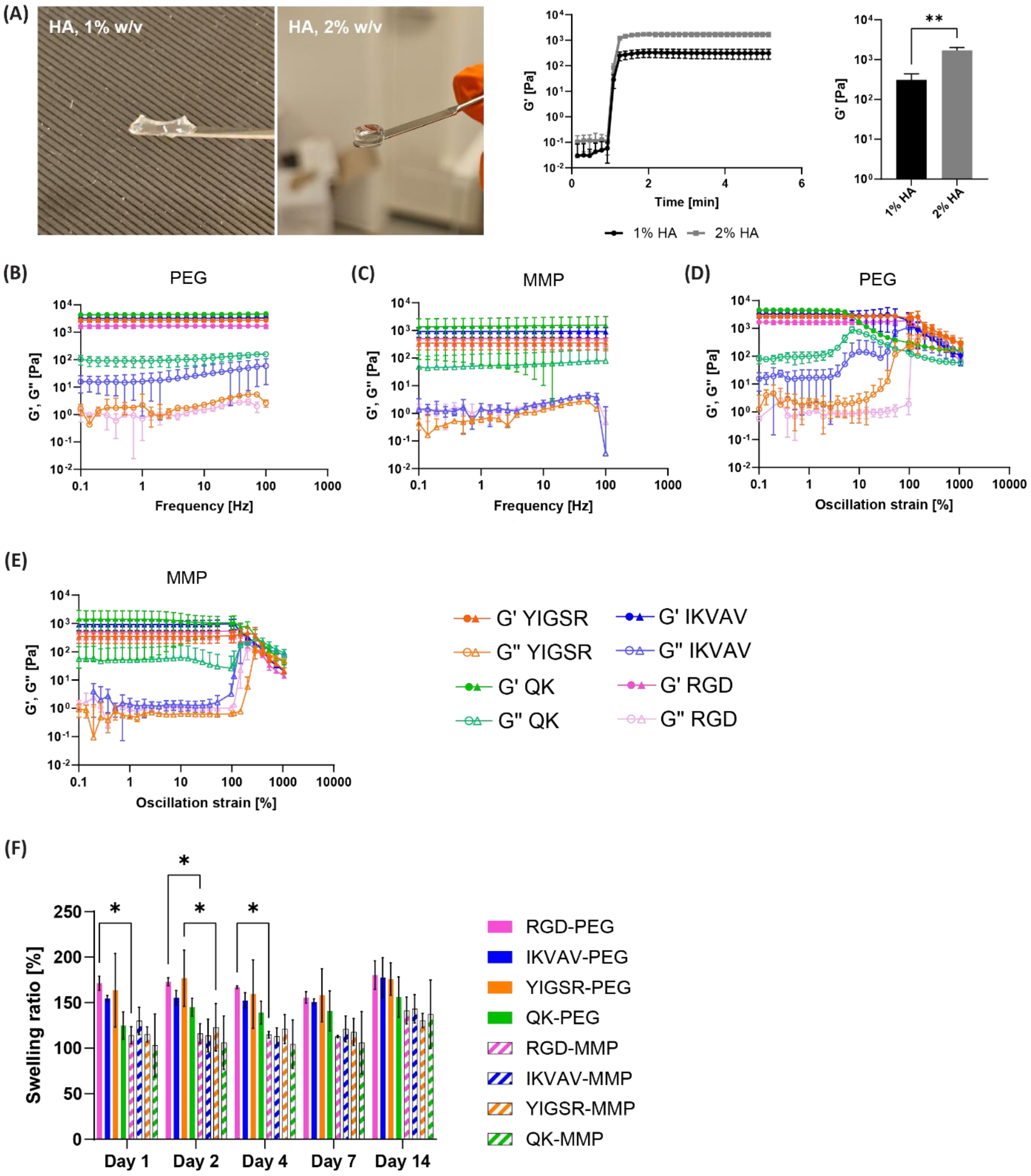


***Figure S3.*** *Characterization of different hydrogel formulations. (A) Qualitative comparison between 1% and 2% w/v hydrogels; rheological time sweep measurement; storage moduli. Frequency sweep step performed on the (B) PEG-crosslinked and (C) MMP-crosslinked hydrogels. Strain sweep step performed on (D) PEG-crosslinked and (E) MMP-crosslinked hydrogel. (F) Water uptake [%] with comparison day by day. Statistical significance represented by: *<0.05.*

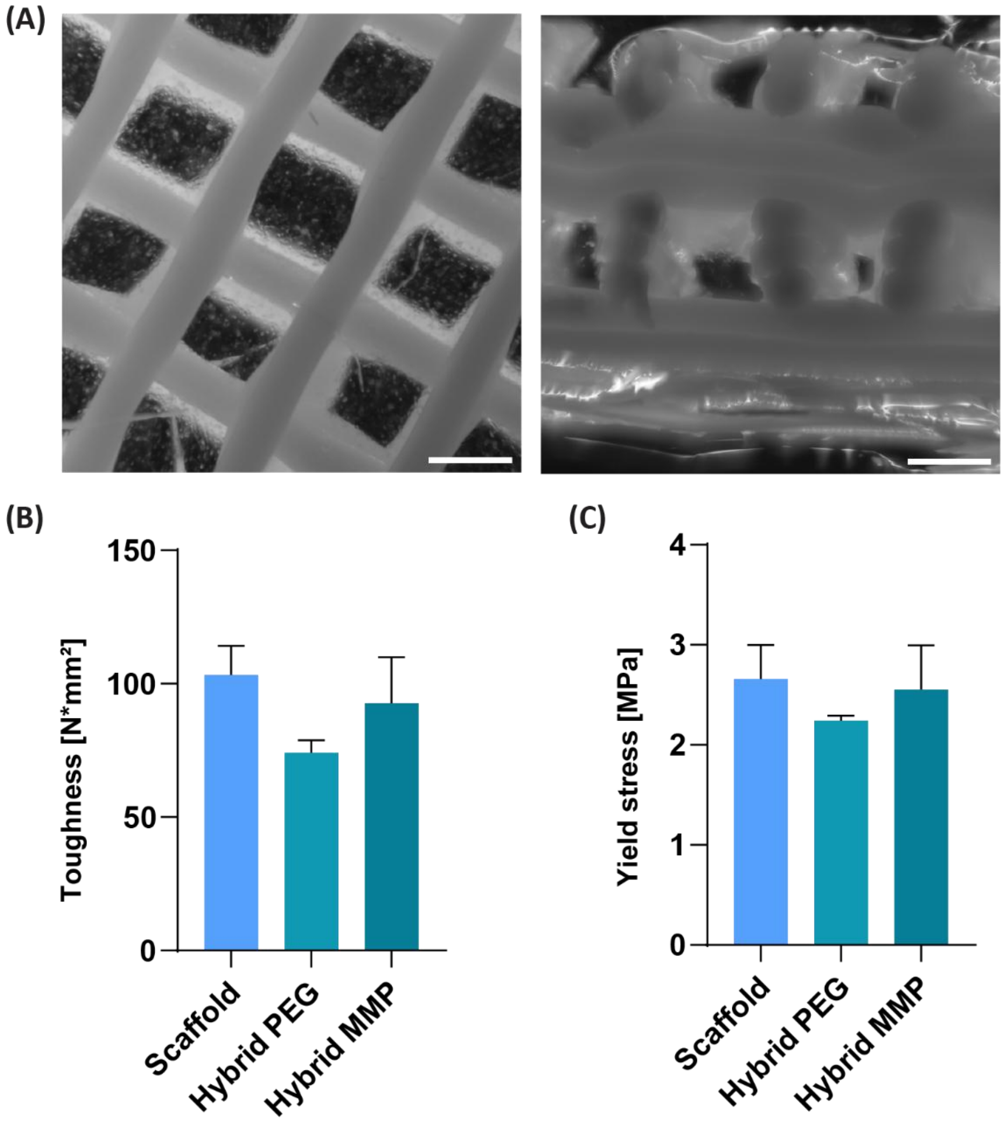


***Figure S4.*** *Characterization of hybrid constructs. (A) Stereomicroscope images of the hybrid constructs, comprising the AM scaffold and the HA hydrogel (top and cross-section view). Scale bar 500 μm. (B) Toughness values extracted from the stress-strain curves showing a comparison among different conditions. (C) Yield stress values extracted from the stress-strain curves showing a comparison among different conditions.*

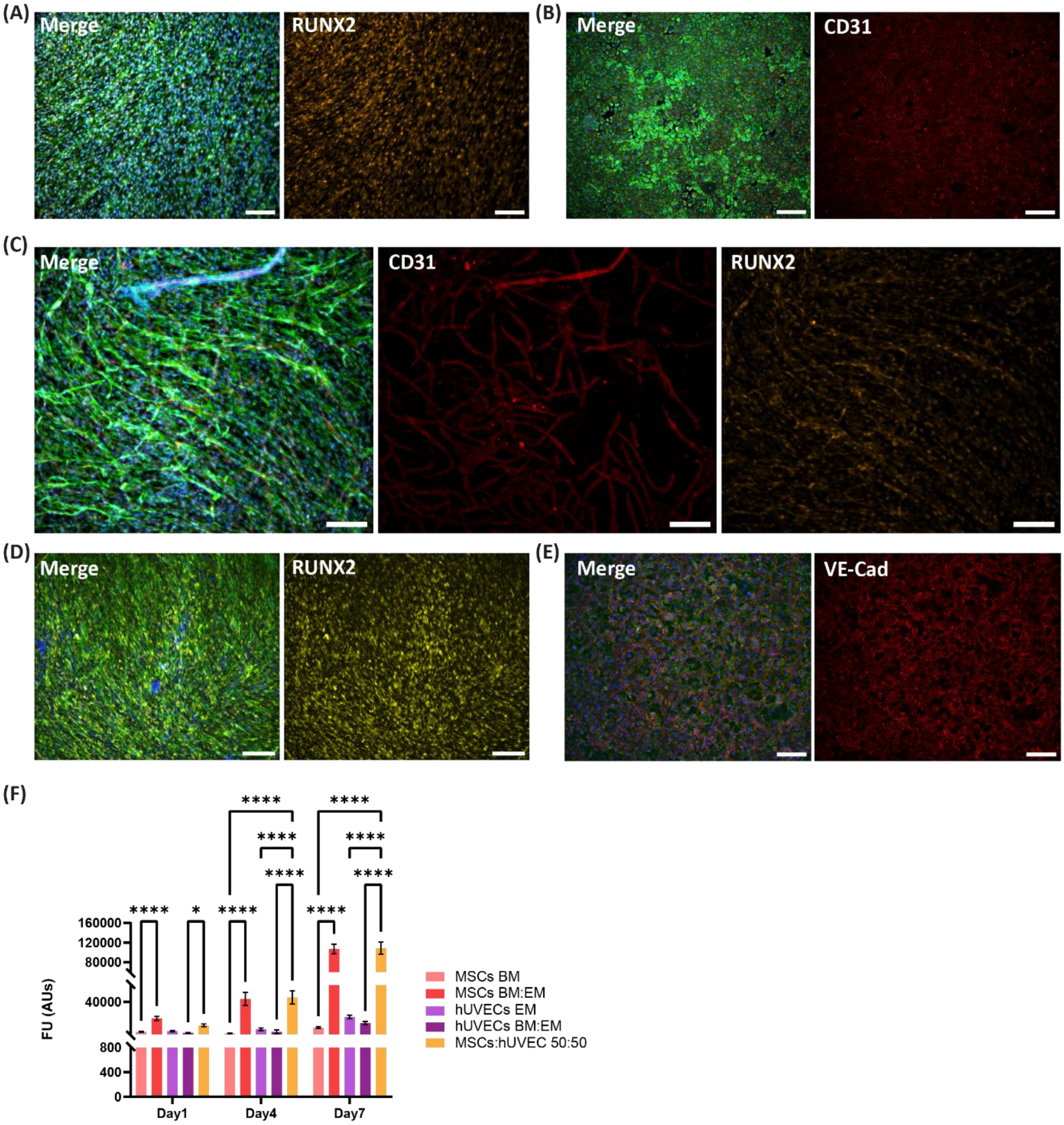


***Figure S5.*** *Immunostaining and metabolic activity of the 2D culture conditions of hMSCs and hUVECs alone or in co-culture, with different media conditions. (A) hMSCs after 7 days in BM:EM 50:50, positive for RUNX2. (B) hUVECs after 7 days in BM:EM 50:50, positive for CD31. (C) hMSC:hUVEC in a 1:1 ratio after 7 days in BM:EM 50:50, showing expression of both RUNX2 and CD31. (D) hMSCs after 7 days in BM only, positive for RUNX2. (E) hUVECs after 7 days in BM only, positive for VE-Cadherin. Scale bar = 200 μm. (F) Metabolic activity measured with Presto Blue assay over 7 days for different culture conditions. Statistical significance represented by: ****<0.0001, *<0.05.*

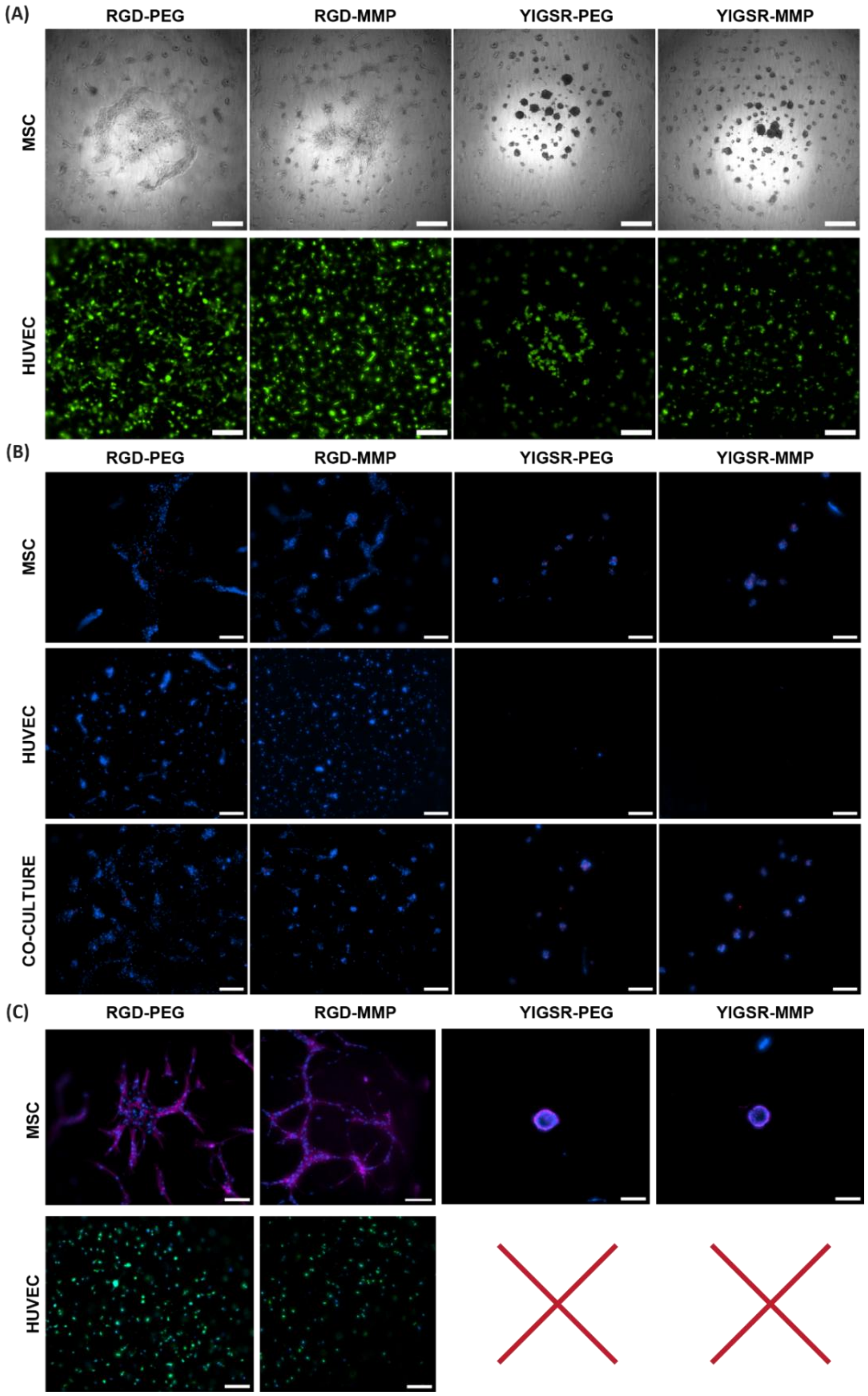


***Figure S6.*** *Assessment of the conditions of cells when seeded on top of the hydrogels. (A) Qualitative evaluation of the morphology of the hMSCs and hUVEC-GFP, cultured alone on each gel, after 24h from the seeding. Scale bar 200 μm. (B) Dapi (blue) and dead (red) staining for all the culture conditions after 24h from the seeding. hUVECs on YIGSR conditions are missing because cells were washed away during staining. Scale bar 200 μm. (C) Immunostaining of the hMSCs and hUVECs after 7 days of culture on different gel conditions. DAPI (blue), Phalloidin (magenta), hUVECs-GFP (green). No hUVECs left on the YIGSR gels after 7 days, due to washing steps involved in the staining procedure. Scale bar 200 μm.*

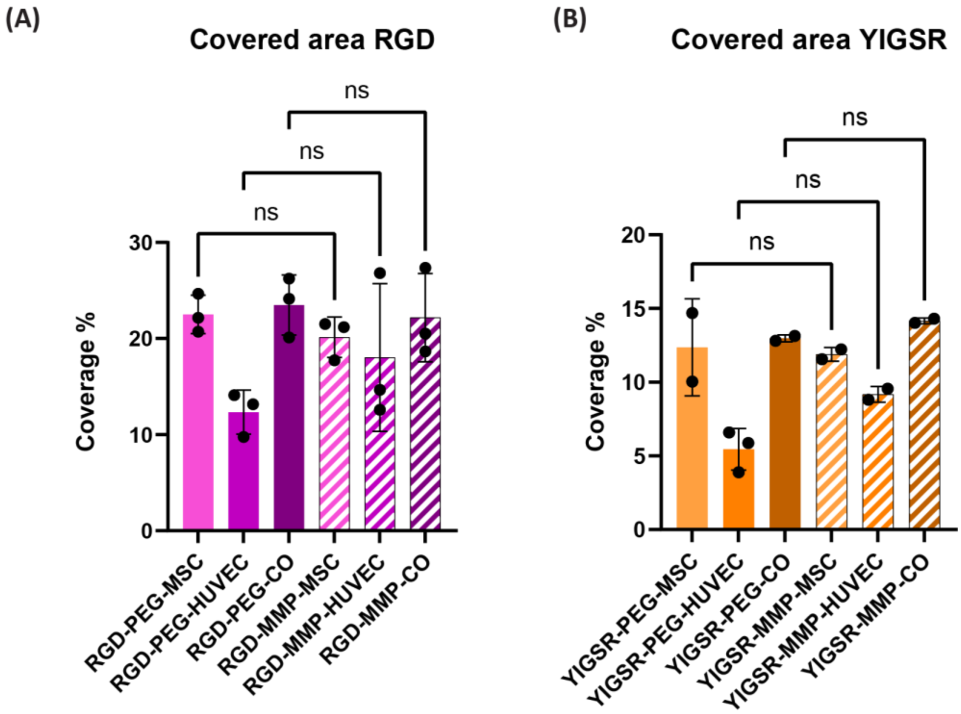


***Figure S7.*** *Assessment of cell response when seeded on top of HA-RGD-PEG/MMP or HA-YIGSR-PEG/MMP hydrogels. Coverage% of the cells seeded on top of the RGD (A) and YIGSR (B) gels. Statistical significance represented by: ns > 0.05.*

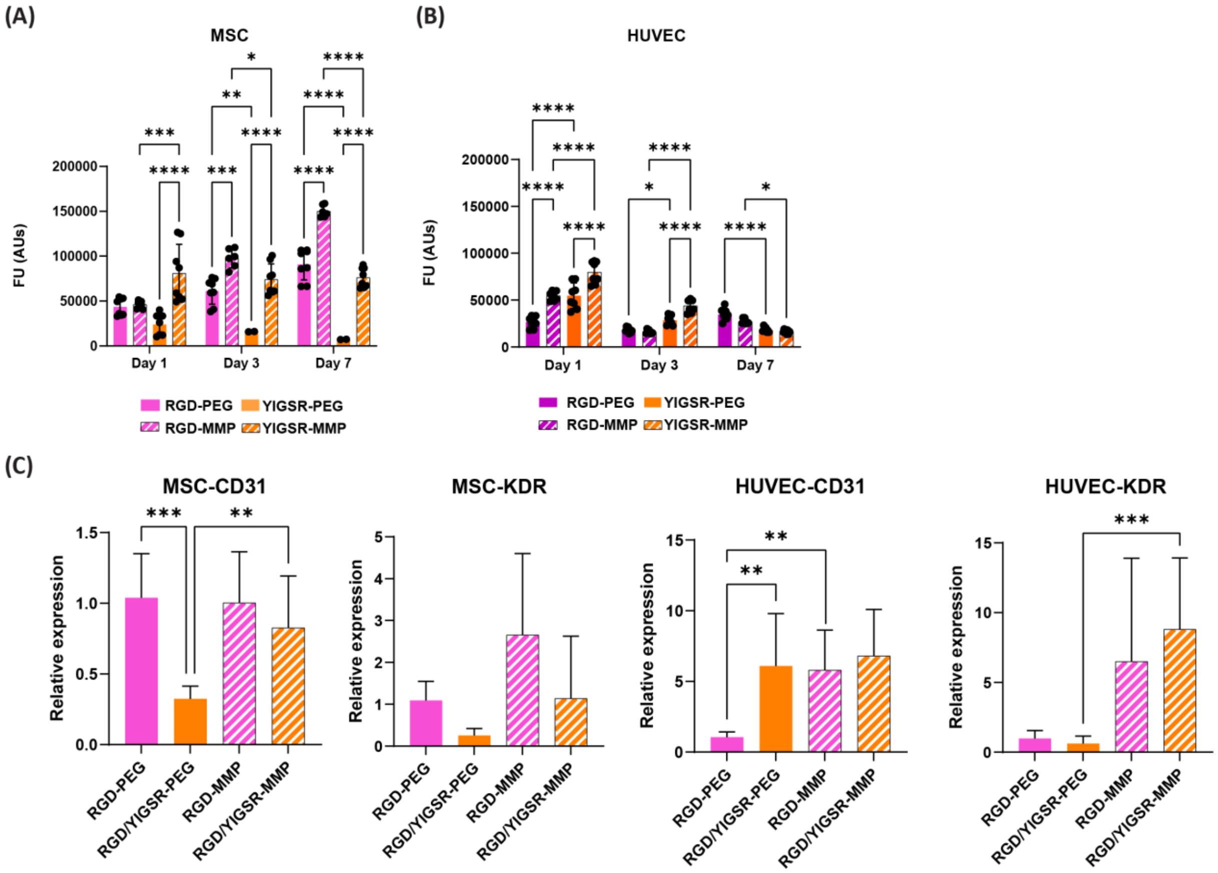


***Figure S8.*** *Assessment of the conditions of cells encapsulated in different hydrogel formulations. Metabolic activity over 7 days of (A) hMSCs and (B) hUVECs. (C) Gene expression of CD31 and KDR genes in the hydrogels only for hMSCs and hUVECs. Values normalized against the RGD-PEG condition. Statistical significance represented by: ****<0.0001, ***<0.001, **<0.01, *<0.05.*

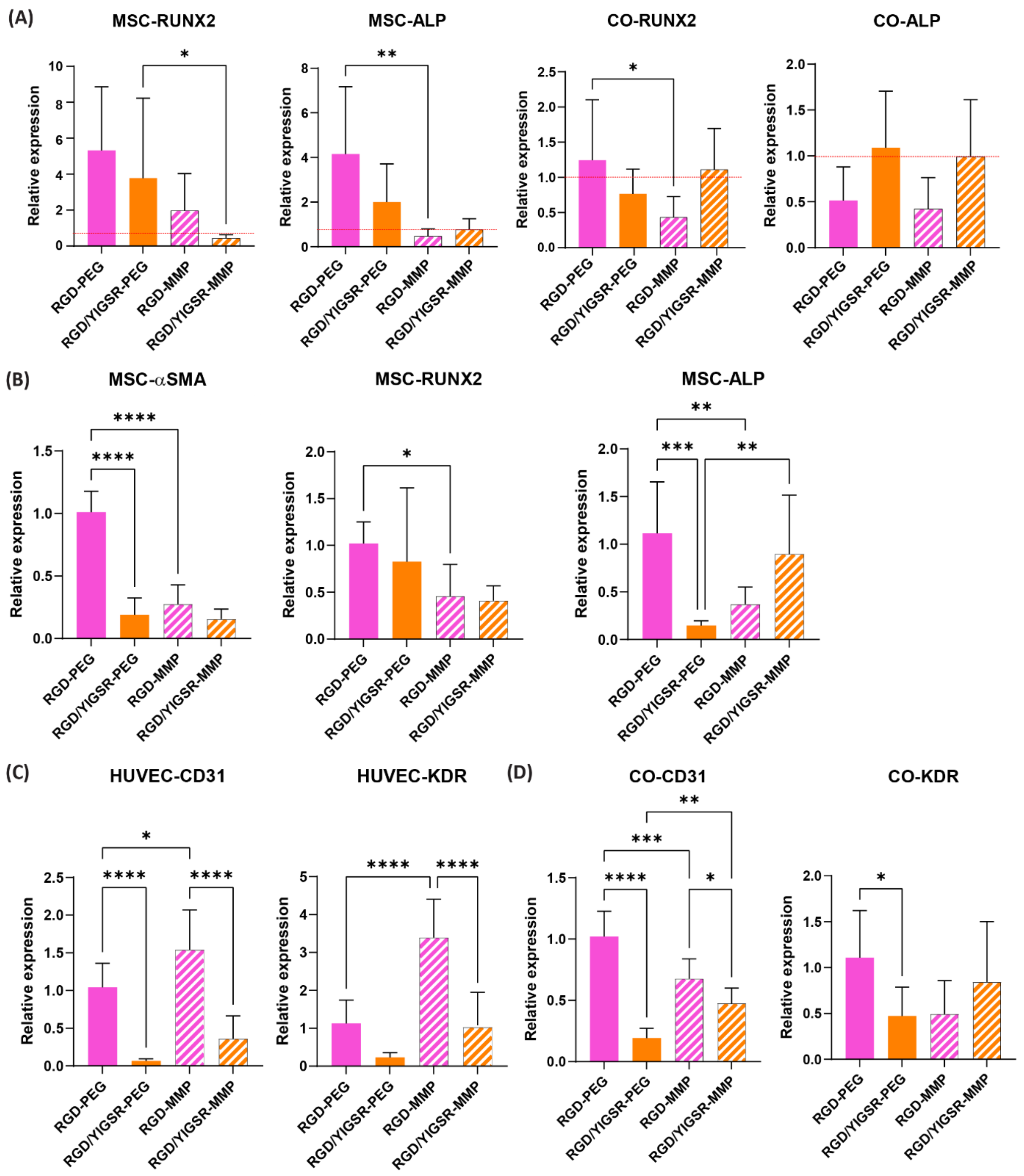


***Figure S9.*** *Assessment of the conditions of cells encapsulated in different hybrid construct formulations. (A) Gene expression of RUNX2 and ALP in the hybrid constructs normalized against the respective hydrogel only conditions, showing that the scaffolds upregulated osteogenic gene expression for hMSCs and co-culture conditions. (B) Gene expression of αSMA, RUNX2 and ALP in hMSCs monocultures in hybrid constructs normalized against the RGD-PEG condition, showing early osteogenic markers in hMSCs. (C) Gene expression of CD31 and KDR in hUVECs monocultures in hybrid constructs normalized against the RGD-PEG condition, showing endothelial markers in hUVECs. (D) Gene expression of CD31 and KDR in the co-culture conditions in hybrid constructs normalized against the RGD-PEG condition, showing endothelial markers in co-culture condition. Statistical significance represented by: ****<0.0001, ***<0.001, **<0.01, *<0.05.*

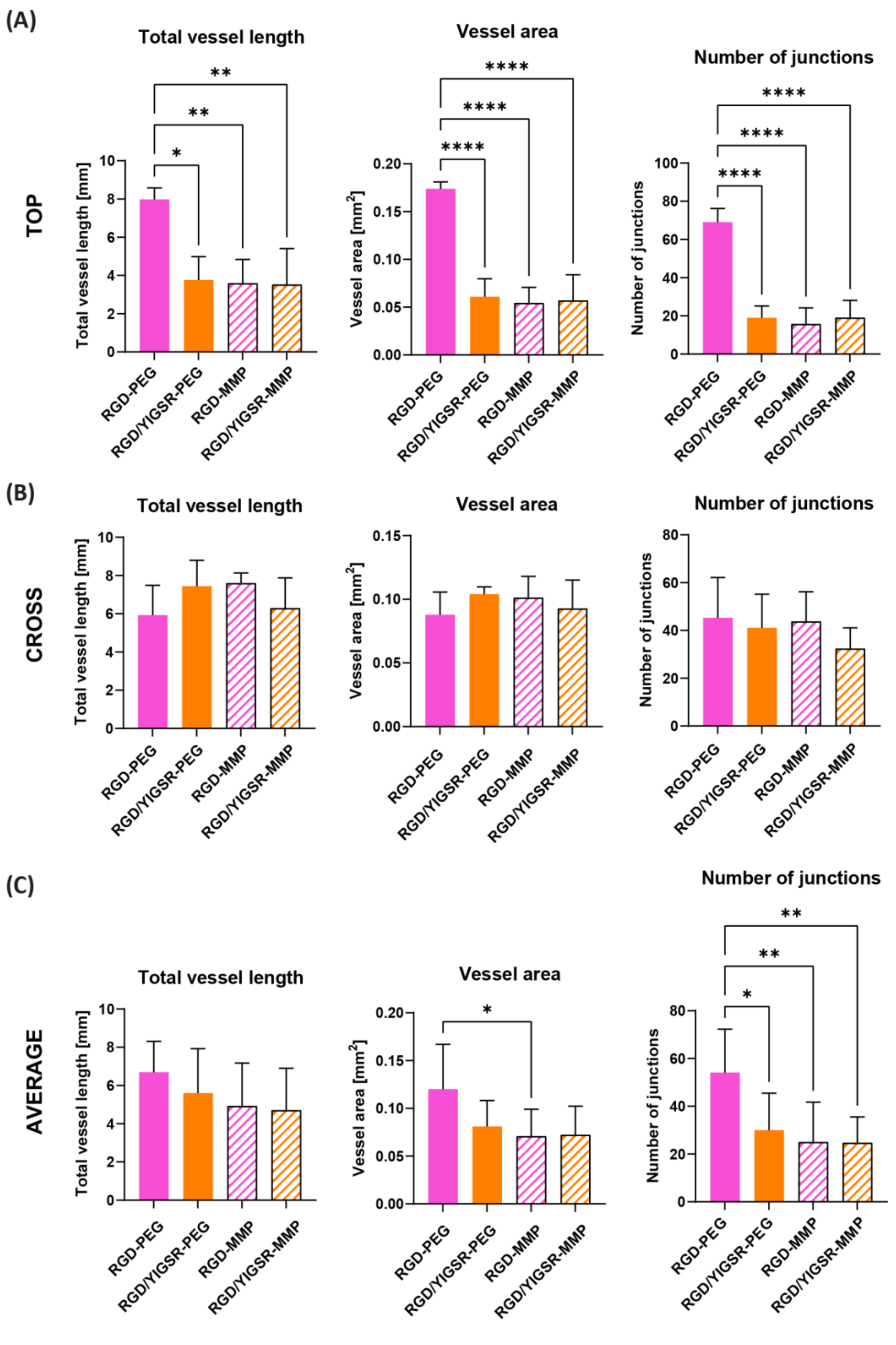


***Figure S10.*** *Quantification of fluorescent images in the co-culture conditions to assess vessel formation parameters after 7 days of culture. Total vessel length, vessel area and number of junctions were quantified on confocal images taken both from the top (A) and from the cross section (B) of at least 3 replicates per condition. An average of top and cross section results was also calculated for a better estimation of the results (C). Statistical significance represented by: ****<0.0001, **<0.01, *<0.05.*
